# Multiviral Quartet Nanocages Elicit Broad Anti-Coronavirus Responses for Proactive Vaccinology

**DOI:** 10.1101/2023.02.24.529520

**Authors:** Rory A. Hills, Tiong Kit Tan, Alexander A. Cohen, Jennifer R. Keeffe, Anthony H. Keeble, Priyanthi N.P. Gnanapragasam, Kaya N. Storm, Michelle L. Hill, Sai Liu, Javier Gilbert-Jaramillo, Madeeha Afzal, Amy Napier, William S. James, Pamela J. Bjorkman, Alain R. Townsend, Mark Howarth

## Abstract

Defending against future pandemics may require vaccine platforms that protect across a range of related pathogens. The presentation of multiple receptor-binding domains (RBDs) from evolutionarily-related viruses on a nanoparticle scaffold elicits a strong antibody response to conserved regions. Here we produce quartets of tandemly-linked RBDs from SARS-like betacoronaviruses coupled to the mi3 nanocage through a SpyTag/SpyCatcher spontaneous reaction. These Quartet Nanocages induce a high level of neutralizing antibodies against several different coronaviruses, including against viruses not represented on the vaccine. In animals primed with SARS-CoV-2 Spike, boost immunizations with Quartet Nanocages increased the strength and breadth of an otherwise narrow immune response. Quartet Nanocages are a strategy with potential to confer heterotypic protection against emergent zoonotic coronavirus pathogens and facilitate proactive pandemic protection.

**One Sentence Summary:** A vaccine candidate with polyprotein antigens displayed on nanocages induces neutralizing antibodies to multiple SARS-like coronaviruses.

## Main

SARS-CoV-2 (SARS2) has caused at least 6 million deaths and new variants continue to emerge (*1*). Despite the success of vaccination in reducing death and serious illness, waning vaccine protection and uncertain efficacy of therapeutics mean that new vaccine strategies are still urgently needed (*2*, *3*). It is also important to protect against new pandemic threats from coronaviruses, which previously led to SARS-CoV (SARS1) and MERS-CoV outbreaks (*4*) and which includes other bat viruses with pandemic potential such as WIV1 and SHC014 (*5*). Immunizing with a single antigen typically induces a narrow strain-specific immune response that may not protect against diverse pre-existing strains or newly arising variants of that pathogen (*6*).

Antigen resurfacing and masking of non-protective regions with glycosylation have been attempted to focus antibody responses on regions of low variability and produce more broadly effective vaccines. However, these strategies can lead to overly specific immune responses susceptible to pathogen escape and have not reliably increased neutralization efficacy (*7*, *8*). Both strategies have been employed on the Receptor-Binding Domain (RBD) of SARS2 Spike. RBD is directly involved in binding to the cell receptor angiotensin-converting enzyme 2 (ACE2) and is the target of most neutralizing antibodies (*9*). Many vaccines have employed RBD as the immunogen and fusion of two SARS2 RBDs into a tandem homodimer was employed early in the COVID-19 pandemic to enhance the immune response, leading to a licensed vaccine (*10*). A tandem heterotrimer composed of one RBD from Wuhan, Beta and Kappa SARS2 has entered clinical development (*11*). Another strategy involves fusion of individual RBDs to proliferating cell nuclear antigen (PCNA) to make a ring with 6 protruding antigens (*12*). However, low molecular weight immunogens may be insufficient to give strong and long-lasting protection (*13*). Highly multivalent display on larger virus-like particles (VLPs) or other nanoparticles enhances the strength and persistence of immune responses, facilitating lymph node uptake and increasing B cell receptor clustering (*13*, *14*). VLP manufacturing uses existing facilities for microbial fermentation to facilitate production of billions of doses (*15*), can avoid the need for a cold-chain (*16*), and has shown a good balance of safety and efficacy (*17*).

In a recently introduced approach, VLPs display a panel of protein variants to favor expansion of B cells recognizing common features of the different antigens. For example, a mosaic of different hemagglutinin heads on ferritin nanoparticles elicited cross-reactive antibodies against diverse influenza strains within the H1 subtype (*18*). This approach has been applied to SARS2, based upon mosaic nanoparticles displaying multiple RBDs from different sarbecoviruses (*6*, *19*, *20*). Sarbecoviruses are the sub-genus of betacoronaviruses that includes SARS1 and SARS2. RBDs can be attached to a VLP through genetic fusion (*20*) or isopeptide coupling (*6*). We previously demonstrated that fusion of a set of sarbecovirus RBDs with SpyTag003 facilitated simple assembly onto the SpyCatcher003-mi3 VLP (*6*) (Fig. 1A). SpyCatcher003 is a protein engineered to rapidly form an isopeptide bond with SpyTag peptide (*21*). mi3 is a 60-mer hollow protein nanocage, computationally designed to self-assemble into a stable dodecahedron (*22*, *23*). In our previous study, the broadest immune response came from mosaic particles displaying 8 different RBDs in a stochastic arrangement (*6*, *19*). These Mosaic-8 nanoparticles elicited neutralizing antibodies against a variety of sarbecoviruses in mouse and rhesus macaque models. Critically, responses were not limited to viruses whose RBDs were represented on Mosaic-8 nanoparticles and included mismatched responses against heterologous sarbecoviruses (*6*, *19*). Mosaic-8 nanoparticles have gained support from the Coalition for Epidemic Preparedness Innovations (CEPI) to enter clinical trials. However, there may be challenges in broad scaling because of the need to produce 9 different components (8 RBDs and SpyCatcher003-mi3) at Good Manufacturing Practice (GMP) level.

**Fig. 1.**
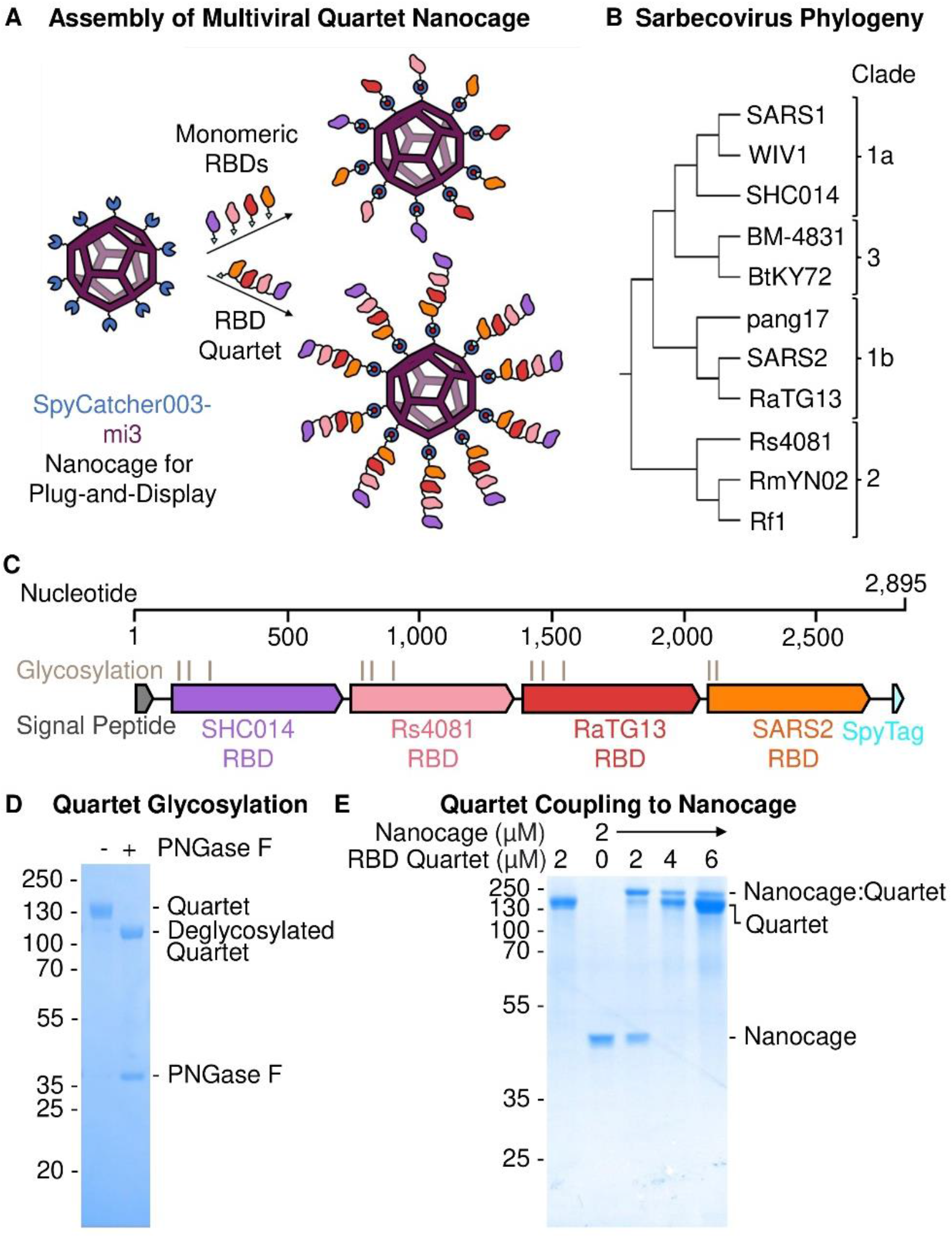
Preparation of Multiviral Quartet. (**A**) Plug-and-Display vaccine assembly of mosaic and Quartet Nanocages. Genetic fusion of SpyCatcher003 (dark blue) with mi3 (purple) allows efficient multimerization of single or Quartet RBDs linked to SpyTag (cyan) through spontaneous isopeptide bond formation (marked in red). Only some antigens are shown in the schematic for clarity. (**B**) Phylogenetic tree of sarbecoviruses used in this study, based on RBD sequence. (**C**) Genetic organization of the multiviral Quartet-SpyTag, indicating the viral origin of RBDs, N-linked glycosylation sites, and tag location. (**D**) Analysis of Quartet-SpyTag with SDS-PAGE/Coomassie staining, with or without PNGase F deglycosylation. (**E**) Coupling of RBD Quartet to SpyCatcher003-mi3 nanocage at different molar nanocage:antigen ratios, analyzed by SDS-PAGE/Coomassie.

Here we establish the production of a multiviral Quartet Nanocage (Fig. 1A). Initially we express a multiviral Quartet from RBDs of 4 different viruses linked as a single polypeptide chain. These antigenic Quartets are assembled via a terminal SpyTag onto SpyCatcher003-mi3 nanocages, creating a protein nanoparticle with dendritic architecture. In addition to reducing the number of vaccine components, this strategy allows a greater number of RBDs to be displayed on each nanocage. We measure antibody responses to the range of sarbecoviruses displayed on the Quartet Nanocage, to sarbecoviruses not present within the chain, as well as to SARS-CoV-2 variants of concern (VOCs). We dissect the breadth of binding to different sarbecoviruses, neutralization potency, and the ability to boost a broad response following focused priming. The magnitude and breadth of antibody induction show that Quartet Nanocages may provide a scalable route to induce neutralizing antibodies across a range of related viruses, to prepare for emerging outbreak disease threats.

### Design of multiviral Quartet Nanocages

RBDs from the evolutionarily-related sarbecoviruses SHC014, Rs4081, RaTG13 and SARS2 Wuhan (Fig. 1B) were genetically fused to produce a multiviral Quartet (Fig. 1C). These RBDs allow comparison to the previously described Mosaic-4 vaccine (*6*). SHC014 can mediate infection of human cells and has been identified as a zoonotic spill-over risk (*24*). Rs4081 is capable of infecting human cells (*25*) but does not enter via ACE2 (*26*). RaTG13 was identified in the intermediate horseshoe bat (*Rhinolophus affinis*) and shares 90% sequence identity for its RBD with SARS2 (Fig. S1 and S2) (*27*). The multiviral Quartet was engineered with a signal sequence for secretion from mammalian cells and a terminal SpyTag, to enable multivalent display on SpyCatcher003-mi3 nanocages (Fig. 1A). The Quartet was secreted efficiently by Expi293F cells and affinity-purified via SpyTag using the SpySwitch system (*28*) (Fig. S3). The Quartet band was relatively broad on SDS-PAGE because of natural variation in glycosylation (Fig. 1D). Removal of N-linked glycans with Peptide N-Glycosidase F (PNGase F) induced a downward shift in protein mobility and a uniform band (Fig. 1D). We demonstrated that the Quartet coupled efficiently to SpyCatcher003-mi3 (Fig. 1E).

### Quartet Nanocages induce antibody responses to diverse sarbecoviruses

We then explored the Quartet’s immunogenicity as a soluble protein or displayed on nanocages, in comparison to a monomeric SARS2 RBD (Fig. 2A). Doses for all immunizations were normalized by the number of SpyTags, allowing comparison of a molar equivalent of SpyCatcher003-mi3 nanocages with similar levels of occupancy. For Uncoupled RBD, this dose of 0.02 nmol corresponds to 0.6 μg protein. Two doses were administered to mice 14 days apart using alum-based adjuvant (Fig. 2B), before quantifying IgG titer against RBD antigens by ELISA. Post-prime, the Quartet Nanocage elicited the highest antibody titer against SARS2 Wuhan RBD, surpassing the Homotypic Nanocage and Uncoupled Quartet (Fig. S4A). Unless indicated, SARS2 responses are measured with Wuhan RBD. We also assessed antibody response to SARS1 RBD, not represented in the immunogens, reflecting induction of broader anti-sarbecovirus antibodies. Here Homotypic Nanocage elicited a weak response against SARS1 but there was still a substantial response from Quartet Nanocage (Fig. S4A). The titer against SARS1 from Quartet Nanocage was greater than the response against SARS2 by Homotypic Nanocage (Fig. S4A).

**Fig. 2.**
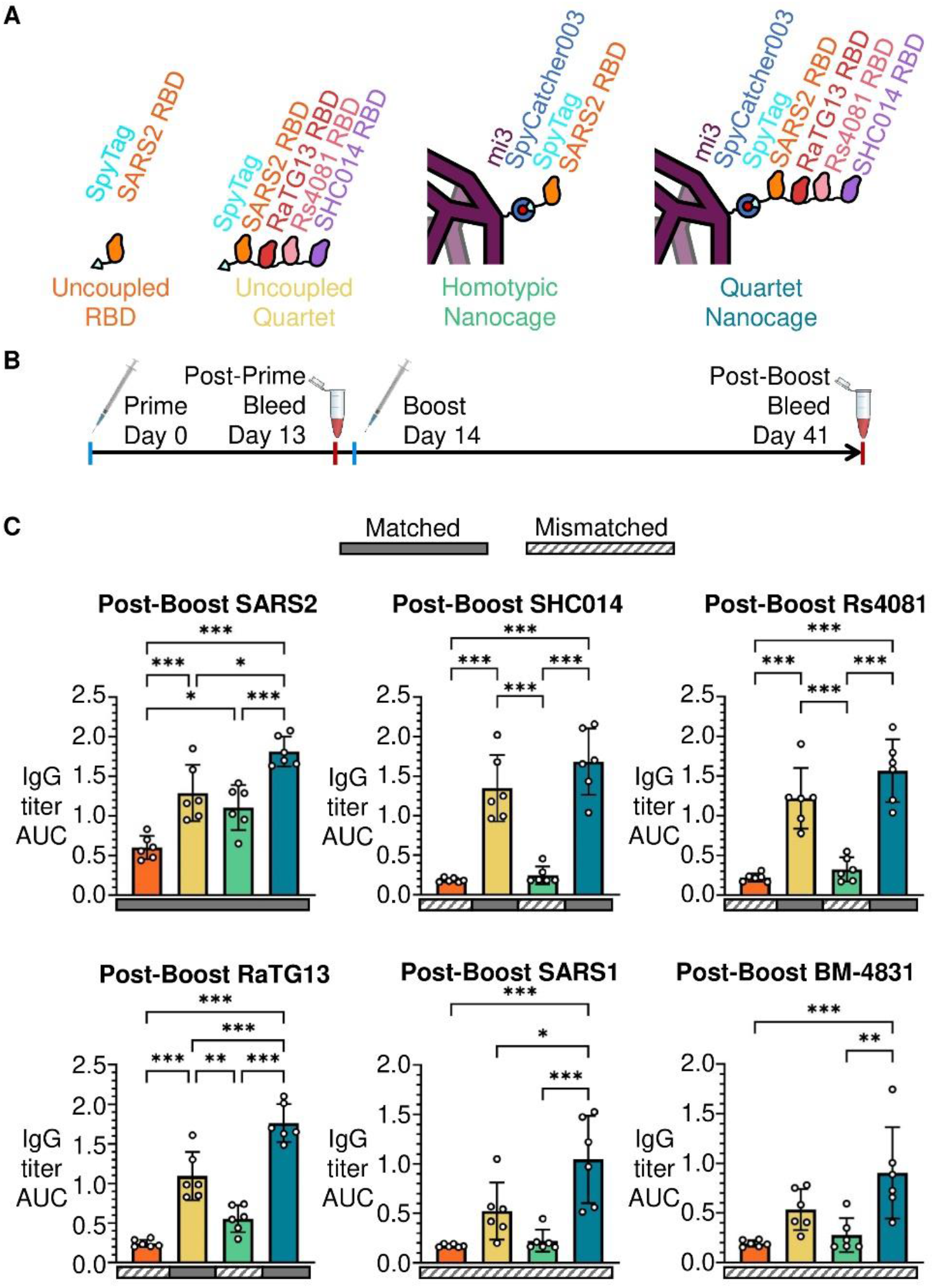
Broad Immune Response from Immunization with Quartet Nanocages. (**A**) Schematic of antigens for this set of immunizations, comparing uncoupled proteins or proteins coupled to the SpyCatcher003-mi3 nanocage. (**B**) Procedure for immunization and sampling. (**C**) ELISA for post-boost serum IgG binding to different sarbecovirus RBDs is shown as the area under the curve (AUC) of a serial dilution. Sera are from mice immunized with uncoupled SARS2 RBD (orange), uncoupled Quartet-SpyTag (yellow), SARS2 RBD coupled to SpyCatcher003-mi3 (green), or Quartet-SpyTag coupled to SpyCatcher003-mi3 (blue). Solid rectangles under samples indicate ELISA against a component of that vaccine (matched). Striped rectangles indicate ELISA against an antigen absent in that vaccine (mismatched). Each dot represents one animal. The mean is denoted by a bar, shown ± 1 s.d., n = 6. * p < 0.05, ** p < 0.01, *** p < 0.001; other comparisons were non-significant.

After boosting, again we found the strongest response against SARS2 from Quartet Nanocage, followed by Uncoupled Quartet, Homotypic Nanocage, and finally Uncoupled RBD (Fig. 2C). This pattern is retained for SARS2 Wuhan, Beta, and Delta Spike (Fig. S4B). After immunizing with Uncoupled RBD or Homotypic Nanocage, responses were very low against other sarbecovirus RBDs (SHC014, Rs4081, RaTG13, SARS1 and BM-4831) (Fig. 2C). However, we saw substantial response against other sarbecovirus RBDs with Uncoupled Quartet and the highest response with Quartet Nanocage (Fig. 2C). We had hypothesized that RBDs exposed at the tip of the Quartet would give stronger responses than RBDs nearer the nanocage surface. In fact, we saw no obvious relationship between the RBD chain location and antibody titer (Fig. 2C). In addition, Quartet Nanocage raised a strong heterotypic response against BM-4831 and SARS1 RBDs absent from the chain, with titers only slightly lower than Homotypic Nanocage against SARS2 (Fig. 2C). Homotypic Nanocage induced its highest cross-reactive response against the closely related RaTG13 RBD, with minimal titers against all other RBDs (Fig. 2C). These results suggest the potential of this Quartet Nanocage approach to induce antibody responses against a broad range of sarbecoviruses.

### Comparison of antibody responses induced by Quartet Nanocages and Mosaic nanoparticles

We next compared the multiviral Quartet to leading mosaic nanoparticle vaccines. Mosaic-4, containing the same 4 RBDs as our Quartet, had induced broad antibodies, but the best breadth was obtained with a Mosaic-8 immunogen (*6*, *19*). Therefore, we also produced the Alternate Multiviral Quartet, containing SpyTag followed by RBDs from other sarbecoviruses: pang17, RmYN02, Rf1 and WIV1 (Fig. S5B). Coupling both the Quartet and Alternate Quartet to SpyCatcher003-mi3 generated the Dual Quartet Nanocage, presenting the same 8 RBDs as Mosaic-8 (Fig. 3A). The Alternate Quartet was efficiently expressed by Expi293F cells and we similarly characterized the protein by SDS-PAGE/Coomassie with and without deglycosylation using PNGase F (Fig. 3B). To interrogate further the relationship between chain position and immunogenicity, we produced a Quartet with SpyTag moved from the C-terminus to the N-terminus (Fig. S5A). This SpyTag-Quartet was used for all subsequent immunizations.

**Fig. 3.**
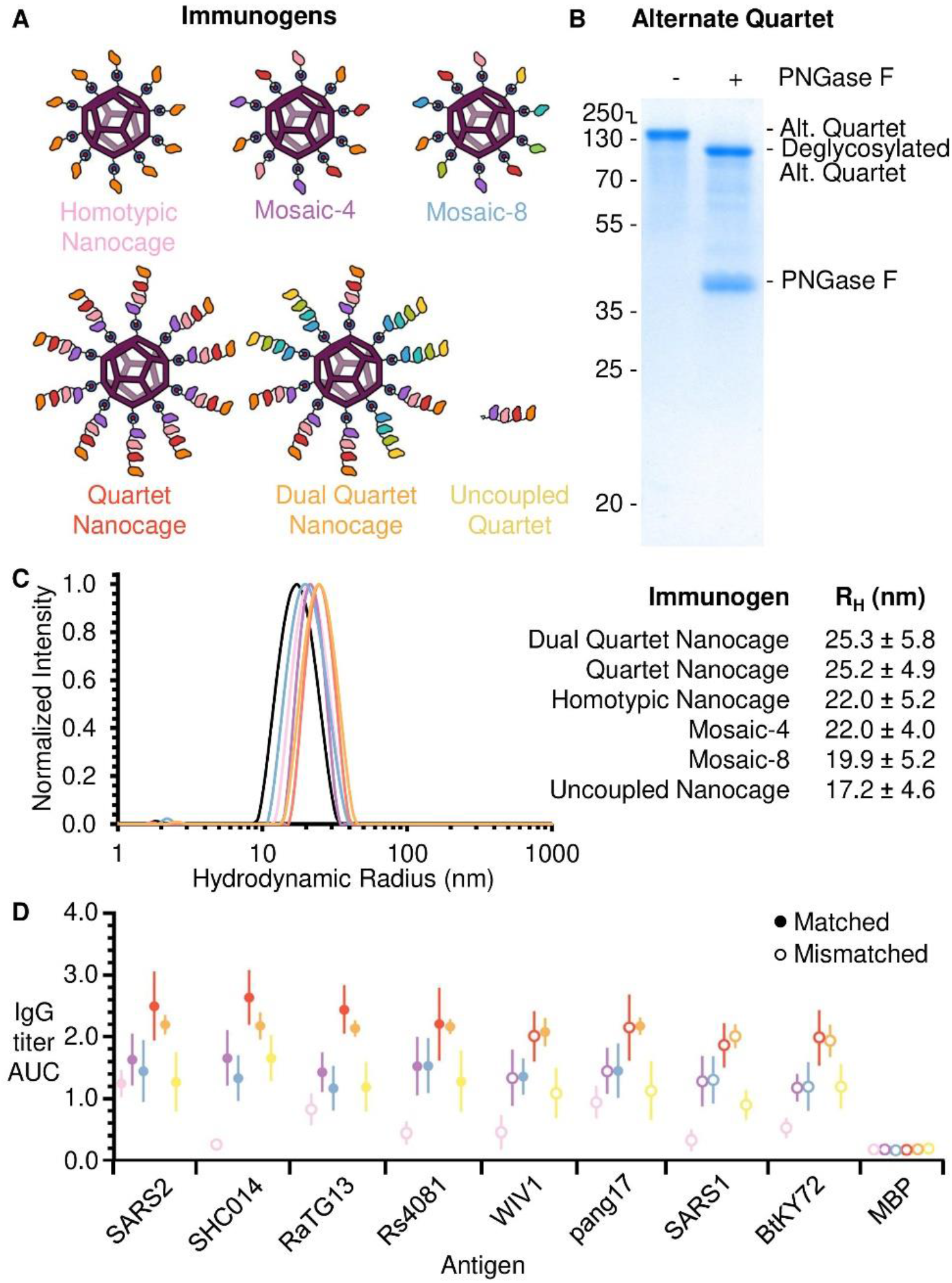
Comparison of immunization with Mosaic or Quartet Nanocages. (**A**) Schematic of antigens for this set of immunizations. (**B**) Validation of the Alternate Quartet by SDS-PAGE with Coomassie staining, shown ± PNGase F deglycosylation. (**C**) DLS of SpyCatcher003-mi3 alone (uncoupled nanocage) or each immunogen. The mean hydrodynamic radius (R_H_) is shown ± 1 s.d., derived from 20 scans of the sample. Uncoupled Nanocage is shown in black, with the other particles colored as in (A). **(D**) ELISA for post-boost serum IgG as the area under the curve of serial dilution, from mice immunized with Homotypic SARS2 Nanocages (pink), Mosaic-4 (purple), Mosaic-8 (blue), SpyTag-Quartet Nanocage (red), Dual Quartet Nanocage (orange), or Uncoupled Quartet (yellow). Filled circles indicate ELISA against a component of that vaccine (matched) while empty circles indicate ELISA against an antigen absent in that vaccine (mismatched). Responses are shown to the set of sarbecovirus RBDs, with SpyTag-MBP as a negative control. The mean is denoted by a circle, shown ± 1 s.d., n = 6. Individual data points and statistics are shown in Fig. S6 and S7.

To compare Mosaic and Quartet Nanocage immunogenicity, we employed a prime-boost approach and analyzed antibody titers, comparing Mosaic-4 and Mosaic-8 with the Quartet Nanocage, Dual Quartet Nanocage, and Uncoupled Quartet (Fig. 3A). Dynamic Light Scattering (DLS) validated that each immunogen homogeneously assembled with SpyCatcher003-mi3 (Fig. 3C). For all RBDs, the two highest post-boost antibody titers were raised by Quartet Nanocage and Dual Quartet Nanocage (Fig. 3D, Fig. S6, Fig. S7). Surprisingly, Quartet Nanocage and Dual Quartet Nanocage induced a similar response to each other against WIV1 and pang17 (Fig. 3D, Fig. S7), even though these antigens were present in Dual Quartet Nanocage but not Quartet Nanocage. In agreement with previous results (*6*), Mosaic-4 and Mosaic-8 produced higher titers than SARS2 Homotypic Nanocage against the RBD set, which were statistically significant with the exception of SARS2. Uncoupled Quartet produced similar titers as both Mosaics against the RBD set, with no statistically significant difference (Fig. 3D, Fig. S6, Fig. S7). These trends were also apparent in post-prime samples, except Mosaic-8 and Quartet Nanocage raised a similar anti-SARS1 response (Fig. S6B). As previously, there was no clear relationship between chain position and antibody response against that RBD. All conditions except Uncoupled Quartet induced a comparable antibody response against SpyCatcher003-mi3 itself (Fig. S6C). SpyTag-Maltose Binding Protein (MBP) was used as a negative control, which revealed minimal antibody response against SpyTag itself (Fig. S6C).

To relate antibody level to antibody efficacy, we tested neutralization of SARS2 Wuhan or Delta virus. We saw the strongest neutralization induced by Quartet Nanocage in each case, while Homotypic Nanocage gave higher responses than Uncoupled Quartet (Fig. 4A, Fig. 4B). To analyze the breadth of neutralizing antibodies, we investigated SARS1 pseudovirus, setting a difficult challenge because SARS1 is a mismatch for all immunogens. Pseudotyped virus neutralization assays correlate well with neutralization of authentic virus (*29*). For this system, we also compared Quartet Nanocage to Mosaic-4 and Mosaic-8. Out of all the immunogens, Dual Quartet Nanocage gave the strongest neutralizing response to SARS1. This was followed by Quartet Nanocage and Mosaic-8, which induced relatively strong and equivalent response against SARS1, while Mosaic-4, Homotypic Nanocage and Uncoupled Quartet gave lower neutralizing responses (Fig. 4C).

**Fig. 4.**
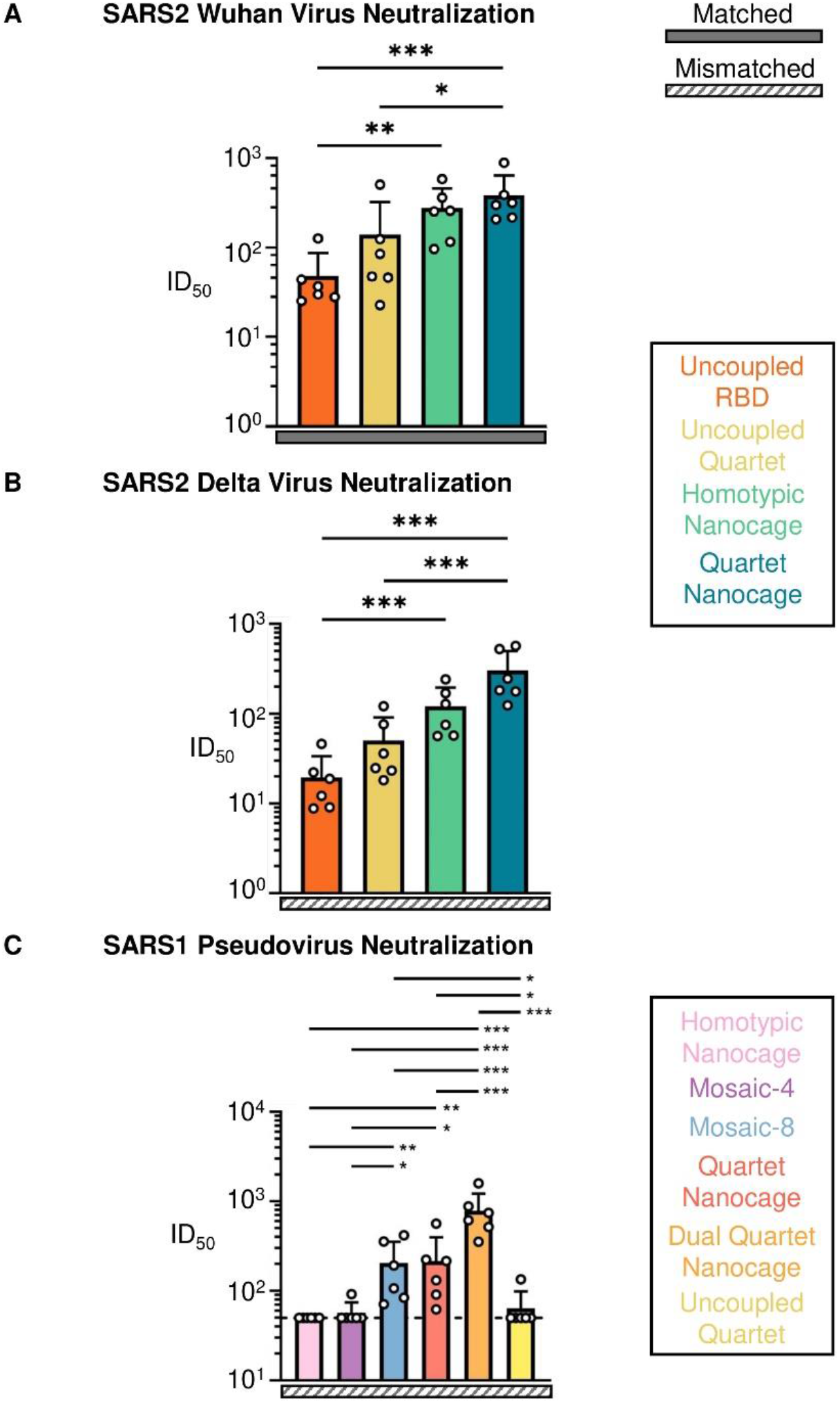
Neutralization induced by Quartet immunogens. (**A**) Neutralization of Wuhan SARS2 virus by boosted mouse sera. Mice were primed and boosted with Uncoupled RBD (orange), Uncoupled Quartet (yellow), Homotypic Nanocage (green), or Quartet Nanocage (blue). Each dot represents one animal, showing the serum dilution giving 50% inhibition of infection (ID_50_). (**B**) Neutralization of Delta SARS2 virus by boosted mouse sera, as in (A). (**C**) Neutralization of SARS1 pseudovirus (mismatched) by post-boost mouse sera, after immunization with different Quartet and Mosaic immunogens. Dashed horizontal lines represent the limit of detection. The mean is denoted by a bar + 1 s.d., n = 6. * p < 0.05, ** p < 0.01, *** p < 0.001; other comparisons were non-significant.

We gained additional insight using 10-fold higher antigen dose and the squalene-based adjuvant AddaVax to enhance viral neutralization further (Fig. S8A). For post-boost sera, there was no significant difference between the antibody titer to SARS2 RBD for any tested vaccine candidate (Fig. S8B). However, the Homotypic Nanocage antibody titer to SARS1 and BtKY72 RBDs were significantly lower than those raised by any of the other conditions, except for the Quartet Nanocage response to SARS1 where the difference did not reach significance (Fig. S8B). Unlike lower-dose immunizations, there was no significant difference between the antibody titer raised by the Mosaic-8, Quartet Nanocage, or Dual Quartet Nanocage to the mismatched SARS1 and BtKY72 RBDs (Fig. S8B). Mosaic-8, Quartet Nanocage, and Dual Quartet Nanocage all elicited favorable neutralization of WIV1, SARS1, and SHC014 pseudovirus (Fig. S8C, Fig. S9). Neutralization of BtKY72 pseudovirus was strong for Mosaic-8 and Dual Quartet Nanocage but less effective for Quartet Nanocage (Fig. S8C). In all cases, Homotypic Nanocage elicited the weakest neutralization of these pseudoviruses (Fig. S8C, Fig. S9A). Under high-dose conditions there was no clear pattern between neutralization by the different immunogens to SARS2 Wuhan, Beta, Delta, and Omicron BA.1 (Fig. S9B, Fig. S10). No immunogen elicited substantial neutralization of SARS2 Omicron XBB.1 pseudovirus (Fig. S9A) or authentic Omicron BQ.1.1 virus using this immunization schedule (Fig. S10), which is consistent with the exceptional immune evasion found for Omicron variants (*30*).

### Quartet Nanocage immunization induces broad antibody responses in animals with a pre-existing focused response

Given the large fraction of the world vaccinated or previously infected with SARS-CoV-2 (671 million confirmed cases and 13 billion vaccine doses administered by February 2023) (*31*, *32*), an outstanding question was whether a broad antibody response could be achieved in the face of a pre-biased immune response. It is not feasible to match the pattern of vaccine sources and timings for different people around the world, but we generated a pre-existing response by priming with SARS2 Wuhan Spike (HexaPro) protein. We then boosted with different immunogens designed to elicit a broad response (Fig. 5A). One hypothesis is that animals with a pre-existing response to SARS2, upon boosting with Quartet Nanocage, would amplify their SARS2 antibodies from a memory response and be less stimulated by other antigens, so the immune response would be narrow. To test this question, we generated Quartet [SARS1], replacing SARS2 with SARS1 RBD (Fig. S5C). This approach led to the ambitious aim of boosting a SARS2 response using an immunogen lacking any SARS2 sequence. We produced Dual Quartet Nanocage [SARS1] by mixing Alternate Quartet and Quartet [SARS1].

**Fig. 5.**
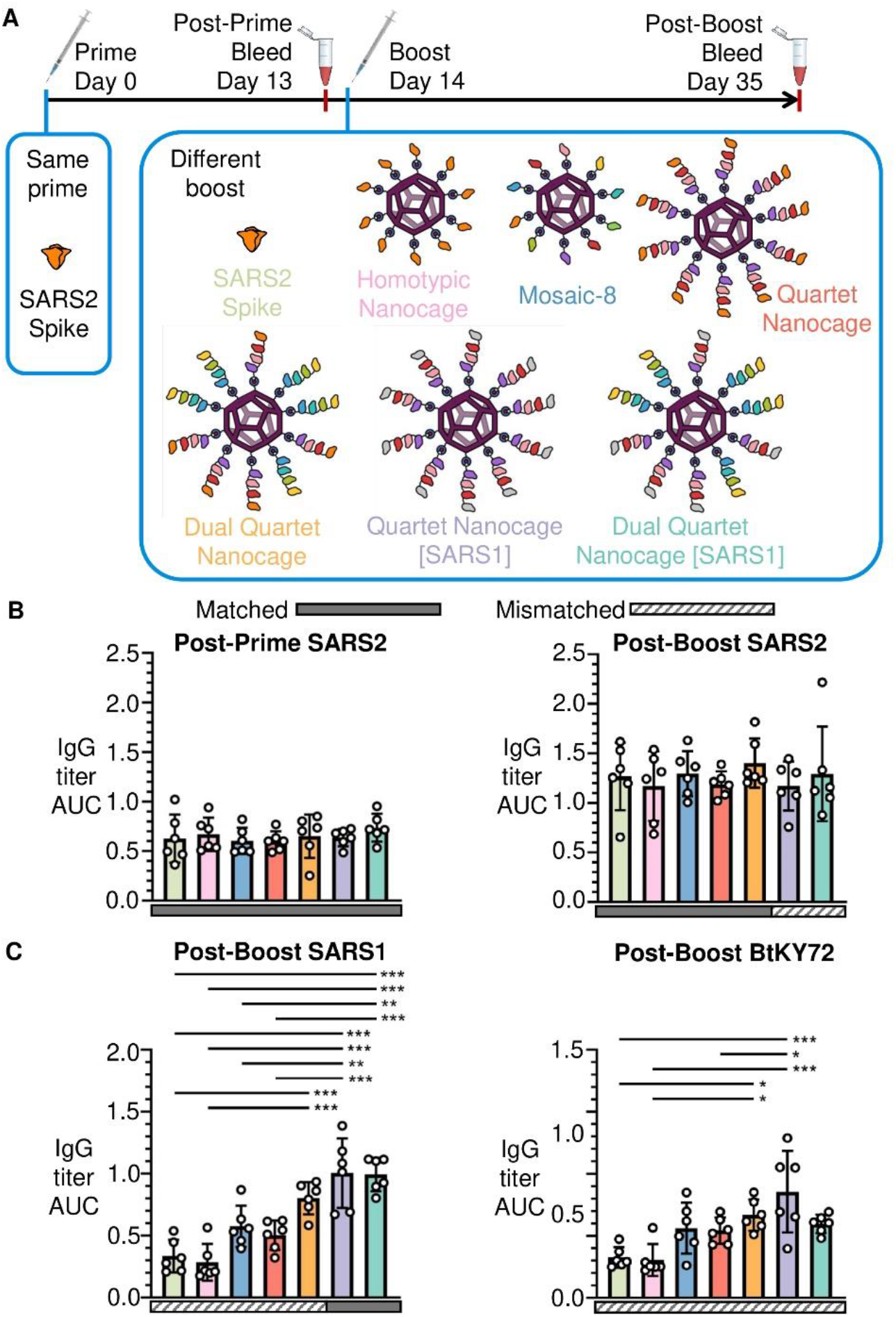
Quartet immunization induces broad antibodies even after a pre-primed SARS2 response. (**A**) Summary of timeline and antigens for this set of immunizations. (**B**) ELISA for serum IgG to SARS2 RBD presented as the area under the curve of a serial dilution. All mice were primed with Wuhan SARS2 Spike, before boosting with Wuhan SARS2 Spike protein (light green), Homotypic Nanocage (pink), Mosaic-8 (dark blue), SpyTag-Quartet Nanocage (red), Dual Quartet Nanocage (orange), Quartet Nanocage with SARS1 RBD replacing SARS2 (purple), or Dual Quartet Nanocage with SARS1 RBD replacing SARS2 (cyan). Solid rectangles under samples indicate ELISA against a component of that vaccine (matched). Striped rectangles indicate ELISA against an antigen absent in that vaccine (mismatched). Each dot represents one animal. The mean is denoted by a bar ± 1 s.d., n = 6. (**C**) ELISA for serum IgG to other sarbecovirus RBDs, as for (B). * p < 0.05, ** p < 0.01, *** p < 0.001; other comparisons were non-significant.

Priming with SARS2 Spike raised the expected narrow strain-specific response against SARS2 RBD (Fig. 5B) and negligible response to SARS1 or BtKY72 (Fig. S11). Surprisingly, the different boosts (Fig. 5B) raised similar responses against SARS2, despite SARS2 RBD being absent in Quartet Nanocage [SARS1] and Dual Quartet Nanocage [SARS1] (Fig. 5B). As expected, Quartet Nanocage [SARS1] and Dual Quartet Nanocage [SARS1] raised the strongest response against SARS1 RBD (Fig. 5C). Quartet Nanocage and Mosaic-8 raised greater antibody response than Homotypic Nanocage or Spike boost against SARS1 and BtKY72 (Fig. 5C). Mismatched responses to SARS1 and BtKY72 raised by Mosaic-8 and Quartet Nanocage were similar to the SARS1 response from a single dose of these candidates in naïve mice (Fig. S6B). Together these results demonstrate that Quartet Nanocages achieve broad anti-sarbecovirus response, despite animals being pre-biased in their response to a specific viral antigen. In addition, Quartet Nanocage lacking SARS2 sequences still induces a good level of anti-SARS2 antibodies, while stimulating broad responses across sarbecoviruses.

## Discussion

Overall, we have established that RBDs from multiple sarbecoviruses can be efficiently expressed as a tandem construct for assembly onto nanocages, creating a dendritic nanoparticle that elicits neutralizing antibodies against SARS2 variants and diverse other sarbecoviruses. Quartet Nanocage enhanced immune response to antigens present on the nanocage, as well as inducing a high level of antibodies to sarbecovirus antigens absent from the particles. Sequential antigen repeats have mostly been explored for strings of T cell epitopes, where there is no folding to a 3D structure or induction of conformation-sensitive antibodies (*33*). Repeats of related structured domains may challenge the cell’s secretion machinery, because of undesired pairings between domains during folding (*34*). However, the cell expression system here efficiently produced the different Quartets that were devised, which may be facilitated by the substantial spacer length and sequence divergence from one domain to the next. In addition, sarbecovirus RBDs exhibit favorable solubility and thermostability (*28*), well suited to advanced antigen assembly strategies.

We were surprised to discover no substantial difference in antibody response to antigens at the start or end of the Quartet. Crystallography or cryoelectron microscopy structures do not allow clear visualization of nanostructures with multiple flexible regions such as the Quartet Nanocage (*35*). Even SpyCatcher003-mi3 coupled to a single SpyTag-RBD showed minimal electron density for RBD in our single-particle cryoelectron microscopy structure (*36*). The glycine/serine linkers between each RBD may provide sufficient flexibility for RBDs near to the nanocage surface to be well exposed to interacting B cells. Upon immunization with Quartet Nanocage, cells with B cell receptors (BCRs) that recognize only a single type of RBD may be less likely to activate efficiently, compared to BCRs recognizing features conserved across sarbecoviruses. Structures have now demonstrated the molecular basis of antibody cross-recognition of diverse sarbecoviruses (*9*, *37*–*47*). Mosaic-8 design was predicated on the idea that stochastic RBD conjugation is ideal for favoring expansion of cross-reactive B cells. However, Mosaic-8 may face challenges in production and regulatory validation. Here the flexibility of the Quartets may achieve a non-uniform surface for B cell stimulation with a uniformly made immunogen. This arrangement also facilitates a greater number of RBDs to be presented per nanoparticle, which may enhance the amount of antibody induction. The vaccine candidates here employ only two (Quartet Nanocage) or three (Dual Quartet Nanocage) components. Despite this, the levels and breadth of antibodies were at least comparable and in many cases higher than the nine component Mosaic-8.

For many diseases, notably malaria and influenza, vaccines face the challenge of inducing novel protective immunity in people with pre-existing immune responses (*48*, *49*). After priming with SARS2 Wuhan Spike, we found that Quartet Nanocages induced an equivalent level of antibodies against Wuhan RBD as more conventional immunogens (Wuhan Spike or Homotypic SARS2 Nanocages). However, Quartet Nanocages additionally broadened response against diverse sarbecovirus RBDs. These data support that a Quartet Nanocage boost could be effective in a human population with existing focused immunity to SARS2.

Limitations of this study are that we immunized only in mice and that expression of tandem antigens was aided by a robust, monomeric antigen; additional optimization would be required for constructing tandem oligomers of obligate trimeric antigens (*50*). There are differences in the vaccine candidates here compared to Mosaic-8b entering clinical trials: here antigens were present on the nanocage at sub-saturating levels with SARS2 Wuhan instead of SARS2 Beta RBD.

We detected antibody induction against the nanocage, but data on VLPs decorated using SpyCatcher or genetic fusion indicate that anti-platform antibodies do not impair responses against the target antigen (*51*, *52*). VLP vaccines have generally shown a good safety margin and scalability for cost-effective global production (*14*, *17*). Nonetheless, in future it may be valuable to apply RBD Quartets using viral vectors (*53*) or mRNA vaccines (*54*) and to explore this approach beyond sarbecoviruses, in other alpha/beta-coronaviruses.

SARS-CoV-2 had a devastating medical and societal impact, despite the rapid generation of effective vaccines. Therefore, it is important that vaccinology possesses further improved tools before the next major viral outbreak (*55*, *56*). The generation of Quartet Nanocages that elicit antibodies across a range of viruses may advance proactive vaccinology, in which broadly-protective vaccines are validated before the pandemic danger emerges (*57*).

## Acknowledgements

We thank Dr. David Staunton from the University of Oxford Department of Biochemistry Biophysical Suite for help with biophysical analysis. We thank the Centre for the AIDS Programme of Research in South Africa (CAPRISA) and Gavin Screaton (University of Oxford) for supplying SARS-CoV-2 variant isolates. The BtKY72 K493Y/T498W Spike plasmid for generating pseudovirus was a kind gift to the Bjorkman lab from David Veesler (University of Washington).

## Funding

Funding for this study was provided by: Biotechnology and Biological Sciences Research Council (BBSRC BB/S007369/1) (A.H.K. and M.H.);

Rhodes Trust (R.H.);

EPA Cephalosporin Early Career Teaching and Research Fellowship (T.K.T.);

Townsend-Jeantet Prize Charitable Trust (Charity Number 1011770) (T.K.T.);

Chinese Academy of Medical Sciences (CAMS) Innovation Fund for Medical Science (CIFMS), China (grant no. 2018-I2M-2-002) (A.R.T.);

University of Oxford COVID-19 Research Response Fund and its donors (reference 0009517) (M.H.) National Institutes of Health (NIH) NIH AI165075 (P.J.B.);

Caltech Merkin Institute (P.J.B.);

George Mason University Fast Grant (P.J.B.).

## Author Contributions

R.A.H. performed all experiments except T.K.T. performed mouse immunizations, J.R.K., P.N.P.G. and K.N.S. tested pseudovirus neutralization, and W.S.J., M.L.H., S.L., J.G-J., M.A. and A.N. tested virus neutralization. A.H.K. designed and purified initial Quartet constructs. R.A.H., T.T., A.A.C., W.S.J., P.J.B., A.R.T. and M.H. designed the project. R.A.H. and M.H. wrote the manuscript. All authors read and approved the manuscript.

## Data availability

Sequences of constructs are available in GenBank, as described in the section “Plasmids and Cloning”. Plasmids encoding pDEST14-SpySwitch, pET28a-SpyCatcher003-mi3, pET28a-SpyTag-MBP, pcDNA3.1-SpyTag-Quartet, pcDNA3.1-Alternate Quartet and pcDNA3.1-Quartet [SARS1] will be deposited before publication in the Addgene repository (https://www.addgene.org/Mark_Howarth/). Further information and request for resources and reagents should be directed to and will be fulfilled by the lead contact, M.H.

## License information

For the purpose of Open Access, the author has applied a CC BY public copyright licence to any Author Accepted Manuscript (AAM) version arising from this submission.

## Supplementary Materials

Materials and Methods

Figs. S1 to S11

