## Supplementary Materials for "Multiviral Quartet Nanocages Elicit Broad Anti-Coronavirus Responses for Proactive Vaccinology"

Materials and Methods
Figs. S1 to S11

### Materials and Methods

#### Plasmids and Cloning

Cloning was performed using standard PCR methods with Q5 High-Fidelity 2× Master Mix (New England Biolabs) and Gibson assembly. All open-reading frames were validated by Sanger sequencing (Source Bioscience).

pET28a-SpyCatcher003-mi3 (GenBank MT945417, Addgene 159995) was previously described (*50*). pET28a-SpyTag-MBP (GenBank MQ038699, Addgene 35050) has been published (*58*). pDEST14-SpySwitch (GenBank ON131074, Addgene plasmid ID 184225) was previously described (*28*). Monomeric sarbecovirus RBD expression vectors contained a C-terminal SpyTag003 (RGVPHIVMVDAYKRYK) (*21*) and $His_8$-tag (*6*) in the plasmid p3BNC-RBD-His8-SpyTag003 and were previously described (*28*): SARS-CoV-2 (GenBank ON131086), SARS-CoV (GenBank ON131087), RaTG13-CoV (GenBank ON131088), SHC014-CoV (GenBank ON131089), Rs4081-CoV (GenBank ON131090), pangolin17 (pang17)-CoV (GenBank ON131091), RmYN02-CoV (GenBank ON131092), Rf1-CoV (GenBank ON131093), WIV1-CoV (GenBank ON131094), Yunnan2011 (Yun11)-CoV (GenBank ON131095), BM-4831-CoV (GenBank ON131096), and BtKY72-CoV (GenBank ON131097)]. The origins of the sarbecovirus RBDs are SARS1 (GenBank AAP13441.1; residues 318-510), WIV1 (GenBank KF367457; residues 307-528), SHC014 (GenBank KC881005; residues 307-524), BM-4831 (GenBank NC014470; residues 310-530), BtKY72 (GenBank KY352407; residues 309-530), pang17 (GenBank QIA48632; residues 317-539), SARS2 (GenBank NC045512; S protein residues 331–529), RaTG13 (GenBank QHR63300; S protein residues 319-541), Rs4081 (GenBank KY417143; S protein residues 310-515), RmYN02 (GSAID EPI_ISL_412977; residues 298-503), and Rf1 (GenBank DQ412042; residues 310-515). The SARS2 Wuhan Spike protein was the HexaPro variant (a gift from Jason McLellan, Addgene plasmid ID 154754) that contains six proline substitutions (F817P, A892P, A899P, A942P, K986P, V987P) that confer greater stability and has been previously described (*59*). The SARS2 Beta variant Spike protein was cloned from HexaPro to match the B.1.351 variant (L18F, D80A, D215G, Δ242-244, R246I, K417N, E484K, N501Y, D614G, A701V) in addition to the previously outlined six proline mutations. The SARS2 Delta variant Spike protein was cloned from HexaPro to match the B.1.617.2 variant (T19R, T95I, G142D, Δ156-157, R158G, L452R, T478K, D614G, P681R, D950N) in addition to the previously outlined six proline mutations.

Quartet RBD constructs were cloned in competent *E. coli* DH5α cells and began with the influenza H7 hemagglutinin (A/HongKong/125/2017) signal-peptide sequence. Each RBD was separated with an 8 or 9 residue Gly-Ser linker that was unique within the construct. pcDNA3.1-Quartet-SpyTag was created by cloning from the N-terminus to C-terminus SHC014 RBD, Rs4081 RBD, RaTG13 RBD and SARS2 RBD with a C-terminal SpyTag into pcDNA3.1 (Fig. 1C, GenBank and Addgene deposition in progress). This is the construct used for Fig. 1 and 2. For subsequent figures, pcDNA3.1-SpyTag-Quartet was cloned with a SpyTag after the signal sequence and then the same order of RBDs (SpyTag-SHC014-Rs4081-RaTG13-SARS2) (Fig. S5, GenBank and Addgene deposition in progress). pcDNA3.1-Quartet [SARS1] was cloned with SpyTag after the signal sequence, with SARS1 in the position of SARS2 (SpyTag-SHC014-Rs4081-RaTG13-SARS1) (Fig. S5, GenBank and Addgene deposition in progress). pcDNA3.1-Alternate Quartet was cloned with SpyTag after the signal sequence, followed by pang17 RBD, RmYN02 RBD, Rf1 RBD, and WIV1 RBD (Fig. S5, GenBank and Addgene deposition in progress).

**Bacterial Expression**

pET28a-SpyCatcher003-mi3 or pET28a-SpyTag-MBP were transformed into *E. coli* BL21(DE3) cells (Agilent) and grown on LB-Agar plates with 50 µg/mL kanamycin for 16 h at 37 °C. A single colony was added in 10 mL LB medium containing 50 µg/mL kanamycin and grown for 16 h at 37 °C with shaking at 200 rpm. This starter culture was then added to 1 L LB containing 50 µg/mL kanamycin and incubated at 37 °C and 200 rpm shaking until $OD_{600}$ 0.6. Cultures were induced with 0.5 mM isopropyl β-D-1-thiogalactopyranoside (IPTG). For SpyCatcher003-mi3, cells were grown at 22 °C with shaking at 200 rpm for 16 h. For SpyTag-MBP, cells were grown at 30 °C with shaking at 200 rpm for 4 h. Cultures were pelleted by centrifugation at 4,000 g.

**Purification of SpyCatcher003-mi3**

Cell pellets were resuspended in 20 mL 20 mM Tris-HCl, 300 mM NaCl, pH 8.5 supplemented with 0.1 mg/mL lysozyme, 1 mg/mL cOmplete mini EDTA-free protease inhibitor (Roche) and 1 mM phenylmethanesulfonyl fluoride (PMSF). The lysate was incubated at 4 °C for 45 min with end-over-end mixing. An Ultrasonic Processor equipped with a microtip (Cole-Parmer) was used to perform sonication on ice (4 times for 60 s, 50% duty-cycle). Centrifugation at 35,000 g for 45 min at 4 °C was used to clear cell debris. 170 mg of ammonium sulfate was added per mL of lysate and incubated at 4 °C for 1 h, while mixing at 120 rpm to precipitate the particles. The solution was centrifuged for 30 min at 30,000 g at 4 °C. The pellet was resuspended in 10 mL mi3 buffer (25 mM Tris–HCl, 150 mM NaCl, pH 8.0) at 4 °C and filtered sequentially through 0.45 µm and 0.22 µm syringe filters (Starlab). The filtrate was dialyzed for 16 h against 1,000-fold excess mi3 buffer. The dialyzed particles were centrifuged at 17,000 g for 30 min at 4 °C and filtered through a 0.22 µm syringe filter. The purified SpyCatcher003-mi3 was loaded onto a HiPrep Sephacryl S-400 HR 16-600 column (GE Healthcare), which was equilibrated with mi3 buffer using an ÄKTA Pure 25 system (GE Healthcare). The proteins were separated at 0.1 mL/min while collecting 1 mL elution factions. The fractions containing the purified particles were pooled and concentrated using a Vivaspin 20 100 kDa molecular weight cut-off centrifugal concentrator (GE Healthcare) and stored at −80 °C.

**Mammalian Protein Expression**

Mammalian expression of all RBD and Spike constructs was performed in Expi293F cells (Thermo Fisher, A14635). Expi293F cells were grown under humidified conditions at 37 °C and 8% (v/v) $CO_2$ in Expi293 Expression Medium (Thermo Fisher) with 50 U/mL penicillin and 50 µg/mL streptomycin. Transfections were performed using the ExpiFectamine 293 Transfection Kit (Thermo Fisher). Expi293F cells were brought to $3\times10^6$ cells/mL and then 1 µg plasmid DNA per mL culture was incubated with ExpiFectamine 293 reagent for 20 min, before being added dropwise to the Expi293F culture. After approximately 20 h, ExpiFectamine 293 Transfection Enhancers 1 and 2 were added. Cell supernatants were harvested after 5 days by centrifuging for 4,000 g at 4 °C for 5 min and were passed through a 0.45 µm filter and then a 0.22 µm filter (Starlab).

**SpySwitch Purification**

RBDs, Quartets and SpyTag-MBP were purified by SpySwitch (*28*). Purifications were performed at 4 °C. For SpyTag-MBP, cells were lysed according to the same procedure as

SpyCatcher003-mi3 and supplemented with 10× SpySwitch buffer (500 mM Tris-HCl pH 7.5 + 3 M NaCl) 10% (v/v). For mammalian proteins, 10 × SpySwitch buffer was added to mammalian culture supernatant at 10% (v/v). SpySwitch resin (*28*), packed in an Econo-Pac Chromatography Column (Bio-Rad), was pre-equilibrated with 2 × 10 column volumes (CV) of SpySwitch buffer (50 mM Tris-HCl pH 7.5 + 300 mM NaCl). The supernatant was incubated with SpySwitch resin for 1 h at 4 °C on an end-over-end rotator. The column was washed twice with 15 CV SpySwitch buffer. Proteins were eluted using a weakly acidic pH switch. The protein was incubated with 1.5 CV of SpySwitch Elution Buffer (50 mM acetic acid/sodium acetate pH 5.0 + 150 mM NaCl) at 4 °C with the column capped. The cap was removed and the elution flow-through was collected into a microcentrifuge tube containing 0.3 CV 1 M Tris-HCl pH 8.0. The microcentrifuge tube was mixed by inversion to minimize the time spent at an acidic pH. This elution step was repeated for a total of six times. Purification was assessed by SDS-PAGE with Coomassie staining. Briefly, 10 µL of fractions were mixed with 2 µL 6× SDS loading buffer [234 mM Tris-HCl pH 6.8, 24% (v/v) glycerol, 120 µM bromophenol blue, 234 mM SDS], before heating at 95 °C for 5 min in a C1000 Touch Thermal Cycler (Bio-Rad) and loading onto 12% SDS-PAGE, before staining with Coomassie. Typical yields for the RBD Quartets are 50-75 mg per L of culture. Typical yields for RBD monomers were 80-160 mg per L of culture, as measured by bicinchoninic acid (BCA). Elution fractions were dialyzed for 16 h against 1,000-fold excess Tris-buffered saline (TBS: 50 mM Tris-HCl, 150 mM NaCl, pH 7.4 at 25 °C). Proteins were stored in aliquots at -80 °C.

**Ni-NTA Purification**
SARS-CoV-2 Spike proteins were purified by nickel-nitrilotriacetic acid (Ni-NTA) affinity chromatography. Mammalian supernatants were supplemented with 10× Ni-NTA buffer (500 mM Tris-HCl, 3 M NaCl, pH 7.8) at 10% (v/v). Ni-NTA agarose (Qiagen) was packed in an Econo-Pac Chromatography Column (Bio-Rad) and washed with 2 × 10 CV of Ni-NTA buffer (50 mM Tris-HCl, 300 mM NaCl, pH 7.8). Mammalian supernatant was incubated in the Ni-NTA column for 1 h at 4 °C with rolling. The supernatant was allowed to flow through by gravity, before being washed with 2 × 10 CV of Ni-NTA wash buffer (10 mM imidazole in Ni-NTA buffer). Elutions were performed by incubating resin with Ni-NTA elution buffer (200 mM imidazole in Ni-NTA buffer) for 5 min, before eluting by gravity. A total of six 1 CV elutions were performed. Elution fractions were assessed by SDS-PAGE with Coomassie staining, pooled, and dialyzed for 16 h against 1,000-fold excess TBS.

**PNGase F Digestion**
Quartet protein (2 µg) was incubated with 1 µL Glycoprotein Denaturing Buffer (10×) (New England Biolabs) at 100 °C for 10 min with a C1000 Touch Thermal Cycler (Bio-Rad). The denatured protein was then chilled on ice for 1 min and centrifuged for 10 s at 2,000 g with a MiniStar silverline (VWR). Then 2 µL GlycoBuffer 2 (10×) (New England Biolabs), 2 µL 10% (v/v) NP-40, 6 µL MilliQ water and 1 µL PNGase F (New England Biolabs) at 500,000 units/mL were added and incubated at 37 °C for 1 h. Proteins were resolved on 12% SDS-PAGE, stained with Coomassie, and imaged using a ChemiDoc XRS imager.

**Dynamic Light Scattering (DLS)**
2 µM SpyTag-antigens were conjugated with 2 µM SpyCatcher003-mi3 for 48 h at 4 °C. Proteins were centrifuged for 30 min at 16,900 g at 4 °C and 30 µL of the supernatant was loaded into a quartz cuvette. Samples were measured at 20 °C using a Viscotek 802 (Viscotek) with 20 scans of 10 s each, using 50% laser intensity, 15% maximum baseline drift and 20% spike tolerance. Before collecting data, the cuvette was incubated in the

instrument for 5 min to allow the sample temperature to stabilize. The intensity of the size distribution was normalized to the peak value using OmniSIZE version 3.0 software, calculating the mean and standard deviation from the multiple scans (Viscotek).

**Endotoxin Depletion and Quantification**

Endotoxin was removed from all vaccine components using Triton X-114 phase separation (*60*, *61*). 1% (v/v) Triton X-114 was added to the protein on ice and incubated for 5 min. The solution was incubated at 37 °C for 5 min and centrifuged for 1 min at 16,000 g at 37 °C. The top phase was transferred to a fresh tube. This procedure was repeated for a total of three times. A final repetition without the addition of Triton X-114 was performed, to account for residual Triton X-114. A Pierce Chromogenic Endotoxin Quant Kit (Thermo Fisher) was used according to manufacturer instructions to quantify the final endotoxin concentration. All vaccine components were below the accepted endotoxin levels for vaccine products of 20 Endotoxin Units (EU) per mL (*62*).

**Immunogen Preparation**

The concentration of vaccine components was measured using BCA assay (Pierce). Where multiple antigens were coupled to the nanocage, the antigens were first mixed in equimolar amounts in TBS. Doses were normalized by the number of SpyTags, to facilitate an equimolar amount of SpyCatcher003-mi3 nanocages with similar occupancy in each condition. For high dose immunizations (Fig. S8-S10), SpyCatcher003-mi3 at 8 µM was incubated with 8 µM SpyTagged antigen for 48 h at 4 °C in TBS pH 8.0. For other immunizations, SpyCatcher003-mi3 at 0.8 µM was incubated with 0.8 µM total SpyTagged antigen for 48 h at 4 °C in TBS, pH 8.0. Uncoupled RBD and Uncoupled Quartet were incubated at 0.8 µM for 48 h at 4 °C in TBS pH 8.0, without the addition of SpyCatcher003-mi3. Prior to immunization, samples were analyzed by SDS-PAGE/Coomassie and DLS. For Fig. 5, SARS2 Spike prime and boost doses were performed with 10 µg SARS2 Wuhan Spike (HexaPro) protein in TBS pH 8.0 at 4 °C.

**Mouse Immunization and Blood Sampling**

Animal experiments were performed according to the UK Animals (Scientific Procedures) Act 1986, under Project License (PBA43A2E4 and PP9362617) and approved by the University of Oxford Animal Welfare and Ethical Review Body. Mice 6 weeks old (at the time of the first immunization) were obtained from Envigo. For high dose immunizations (Fig. S8-S10), we used BALB/c female mice and for all other immunizations we used C57BL/6 female mice. Mice were housed in accordance with the UK Home Office ethical and welfare guidelines and fed on standard chow and water *ad libitum*. Prior to immunization, immunogens were mixed 1:1 with VAC 20 adjuvant (SPI Pharma) (25 µL + 25 µL), except for the high dose immunizations (Fig. S8-S10) where immunogens were mixed 1:1 with AddaVax (Invivogen). This procedure gave final doses of 0.2 nmol total SpyTagged antigen for high dose immunizations and 0.02 nmol total SpyTagged antigen for normal dose immunization. For normal dose immunization, this relates to 0.6 µg Uncoupled RBD. Isoflurane (Abbott)-anesthetized mice were immunized on day 0 and day 14 intramuscularly in the gastrocnemius muscle with the specified antigen-adjuvant mix. Post-prime blood samples were obtained on day 13 via tail vein using Microvette (CB300, Sarstedt) capillary tubes. Post-boost samples were obtained on Day 32 to 41 (exact day for each set of immunizations is indicated in the figure) via cardiac puncture of humanely sacrificed mice. The collected whole blood in microtainer SST tubes (Becton Dickinson) was allowed to clot at 25 °C for 1-2 h, before spinning down at 10,000 g for 5 min at 25 °C. The sera were heat-inactivated at 56 °C for 30 min, before storing at −20 °C.

**Enzyme-Linked Immunosorbent Assay (ELISA)**

Nunc MaxiSorp plates (Thermo Fisher) were coated with 80 nM purified SpyTag-RBD, SpyTag-MBP or SpyCatcher003-mi3 in PBS (137 mM NaCl, 2.7 mM KCl, 10 mM $Na_2HPO_4$, 1.7 mM $KH_2PO_4$, pH 7.4) at 4 °C for 16 h. Where SARS2 was analyzed, this refers to the Wuhan variant, unless indicated. In Fig. S4B, the response to different SARS2 variants was measured by coating 1 µg/mL of the indicated HexaPro Spike protein in PBS and incubating at 4 °C for 16 h. Plates were washed three times with PBS supplemented with 1% (v/v) Tween 20 (PBST). Plates were blocked by 2 h incubation at 25 °C with 5% (w/v) skimmed milk in PBS. Plates were then washed three times with PBST. Sera were serially diluted into the blocking buffer using 8-point, 4-fold series starting at 1:100. Plates were incubated with sera for 1 h at 25 °C and then washed three times with PBST. Plates were incubated at 25 °C for 1 h with a 1:1,600 dilution of horseradish peroxidase-conjugated goat anti-mouse IgG antibody (Sigma-Aldrich A9044). Plates were washed three times with PBST. Plates were then incubated at 25 °C for 5 min with 1-Step™ Ultra TMB-ELISA Substrate Solution (Thermo Scientific) before the reaction was stopped with 1 M $H_2SO_4$. $A_{405}$ measurements were taken with a FLUOstar Omega plate reader (BMG Labtech) using Omega MARS software (BMG Labtech). A sigmoidal dose response curve was fit to the absorbance data using the optimize.curve_fit() function from the Python SciPy library (*63*). The sigmoidal dose response function was:

$$\mathrm{y} = \mathrm{Bottom} + \frac{\mathrm{Top} - \mathrm{Bottom}}{1 + 10^{\log_{10}(\mathrm{IC}_{50}) - \mathrm{x}}}$$

The area under the fitted curve (AUC) was determined using the trapz function from the Python Numpy library (*64*). Area under the curve was used instead of endpoint titer to account better for data across the entire range of values (*65*). Results were plotted using GraphPad Prism (GraphPad Software version 9.4.1).

**Microneutralization Assay**

These assays were performed in the James & Lillian Martin Centre, University of Oxford, operating under license from the Health and Safety Authority, UK, on the basis of an agreed Code of Practice, Risk Assessments (under the Advisory Committee on Dangerous Pathogens) and standard operating procedures. The microneutralization assay determines the serum concentration that induces a 50% reduction in focus-forming units of SARS2 in Vero cells (American Type Culture Collection, CCL-81). A serial dilution of immunization sera [seven steps from 1/40 to 1/40,000 diluted into Dulbecco's Modified Eagle Medium (DMEM)] was pre-incubated for 30 min at 25 °C with a fixed dose of 100-200 focus-forming units (20 µL) of different authentic SARS-CoV-2 variants. This procedure was performed in triplicate for samples from high dose immunizations outlined in Fig. S8-S10 and in quadruplicate for all other samples. DMEM on its own was used for serum-free control wells, which were used to define 100% infectivity. The Victoria 01/2020 isolate (Pango B) was used for Wuhan neutralization (*66*). The Beta variant (Pango B.1.351) used for neutralizations is the HV001 isolate, sequenced and kindly supplied by CAPRISA, Durban, South Africa (*67*). The isolates for Delta (Pango B.1.617.2), Omicron BA.1 (Pango B.1.1.529.1), and Omicron BQ.1.1 (Pango B.1.1.529.5.3.1.1.1.1.1.1) were kindly supplied by Gavin Screaton (University of Oxford). This mixture was incubated with 100 µL of Vero cells ($4.5 \times 10^4$) at 37 °C with 5% (v/v) $CO_2$. 2 h into this incubation, a 1.5% (w/v) carboxymethyl cellulose-containing overlay was applied in order to prevent satellite focus formation. 18 h post-infection, the monolayers were fixed with 4% (w/v) paraformaldehyde in PBS and then permeabilized with 2% (v/v) Triton X-100. The cells were stained using the FB9B monoclonal antibody at 1 µg/mL (*68*). These samples were developed using an anti-human IgG (Fc-specific) peroxidase-conjugated antibody (1:5,000 dilution, cat. no. A0170-1ML, Sigma-Aldrich) and True Blue peroxidase substrate. The infectious foci were enumerated by Classic ELISpot Reader (AID GmbH). Data were analyzed using four-parameter logistic regression (Hill equation) using GraphPad Prism (GraphPad Software version 8.3). Statistical significance of differences between groups was determined using a one-way analysis of variance (ANOVA) test, followed by Tukey's multiple comparison post hoc test of $ID_{50}$ values converted to $\log_{10}$ scale using GraphPad Prism (GraphPad Software version 9.4.1).

**Pseudovirus Neutralization Assay**

SARS2 BQ.1.1, SARS1, WIV1, SHC014, and BtKY72 K493Y/T498W pseudotyped viruses were prepared as described (*69*, *70*). The double mutation BtKY72 K493Y/T498W in the BtKY72 Spike protein has previously been shown to enable entry to human cells via ACE2 (*26*). This technique for producing pseudoviruses employs HIV-based lentiviral particles with genes encoding the appropriate Spike protein lacking the cytoplasmic tail. A three-fold serial dilution of sera was incubated with pseudotyped virus for 1 h at 37 ˚C. The mixture was incubated with $293T_{ACE2}$ target cells for 48 h at 37 ˚C (*6*). Cells were washed twice with PBS, before being lysed with Luciferase Cell Culture Lysis 5× reagent (Promega). NanoLuc Luciferase activity in the lysates was measured using the Nano-Glo Luciferase Assay System (Promega). The relative luminescence units (RLUs) were normalized to values derived from cells infected with pseudotyped virus in the absence of serum. Half-maximal inhibitory dilution ($ID_{50}$) was determined using 4-parameter nonlinear regression in AntibodyDatabase (*71*) and plotted using using GraphPad Prism (GraphPad Software version 9.4.1). Statistical significance of differences between groups was determined using an ANOVA test, followed by Tukey's multiple comparison post hoc test of $ID_{50}$ values converted to $\log_{10}$ scale using GraphPad Prism (GraphPad Software version 9.4.1).

**Bioinformatics**

The phylogenetic tree of sarbecovirus RBD sequences was constructed using MEGA X v 11.0.13 software (*72*). Multiple sequence alignment and calculation of amino acid identity was performed using Clustal Omega v 1.2.4 (*73*). The structure of SARS2 RBD was based on PDB ID: 6ZER (*74*) and analyzed using PyMOL version 2.5.2.

**Statistics and Reproducibility**

No statistical method was used to predetermine sample size. Significance for ELISAs was calculated with an ANOVA test using Tukey's post hoc test in GraphPad Prism (GraphPad Software version 9.4.1). Comparisons for neutralizations were calculated with an ANOVA test, followed by Tukey's multiple comparison post hoc test of $ID_{50}$ values converted to $\log_{10}$ scale using GraphPad Prism (GraphPad Software version 9.4.1). Stars were assigned according to: * $p < 0.05$, ** $p < 0.01$, *** $p < 0.001$. On graphs where some conditions are compared, where no test is marked then the difference was non-significant. The experiments were not randomized. The Investigators were not blinded to allocation during experiments and outcome assessment.

```
        320                 340                 360                 380                 400
Rs4081  RVSPTHEVVRFPNITNRCPFDKVFNASRFPNVYAWERTKISDCVADYTVLYNS-TSFSTFKCYGVSPSKLIDLCFTSVYADTFL
RmYN02  RILPSTEVVRFPNITNFCPFDKVFNATRFPNVYAWQRTKISDCIADYTVLYNS-TSFSTFKCYGVSPSKLIDLCFTSVYADTFL
Rf1     RVSPVTEVVRFPNITNLCPFDKVFNATRFPSVYAWERTKISDCVADYTVFYNS-TSFSTFNCYGVSPSKLIDLCFTSVYADTFL
BM-4831 RVTPTTEVVRFPNITQLCPFNEVFNITSFPSVYAWERMRITNCVADYSVLYNSSASFSTFQCYGVSPTKLNDLCFSSVYADYFV
BtKY72  RVSPSTEVVRFPNITNLCPFGQVFNASNFPSVYAWERLRISDCVADYAVLYNSSSSFSTFKCYGVSPTKLNDLCFSSVYADYFV
pang17  RVQPTISIVRFPNITNLCPFGEVFNASKFASVYAWNRKRISNCVADYSVLYNS-TSFSTFKCYGVSPTKLNDLCFTNVYADSFV
SARS2   RVQPTESIVRFPNITNLCPFGEVFNATRFASVYAWNRKRISNCVADYSVLYNS-ASFSTFKCYGVSPTKLNDLCFTNVYADSFV
RaTG13  RVQPTDSIVRFPNITNLCPFGEVFNATTFASVYAWNRKRISNCVADYSVLYNS-TSFSTFKCYGVSPTKLNDLCFTNVYADSFV
SHC014  RVAPSKEVVRFPNITNLCPFGEVFNATTFPSVYAWERKRISNCVADYSVLYNS-TSFSTFKCYGVSATKLNDLCFSNVYADSFV
SARS1   RVVPSGDVVRFPNITNLCPFGEVFNATKFPSVYAWERKKISNCVADYSVLYNS-TFFSTFKCYGVSATKLNDLCFSNVYADSFV
WIV1    RVAPSKEVVRFPNITNLCPFGEVFNATTFPSVYAWERKRISNCVADYSVLYNS-TSFSTFKCYGVSATKLNDLCFSNVYADSFV
        *: *  .::******: ***..*** : * .****:* :*::*:***:*:*** : ****:***** :** ****:.**** *:

                         420                 440                 460                 480
Rs4081  IRSSEVRQVAPGETGVIADYNYKLPDDFTGCVIAWNTAKQDQG-----QYYYRSSRKTKLKPFERDLTSDE-------------
RmYN02  IRFSEVRQIAPGETGVIADYNYKLPDDFTGCVLAWNTAQQDIG-----SYFYRSHRAVKLKPFERDLSSDE-------------
Rf1     IRFSEVRQVAPGQTGVIADYNYKLPDDFTGCVIAWNTAKQDVG-----SYFYRSHRSSKLKPFERDLSSEE-------------
BM-4831 VKGDDVRQIAPAQTGVIADYNYKLPDDFTGCVIAWNTNSLDS--SNE--FFYRRFRHGKIKPYGRDLSNVLFNPSGGTCSA-EG
BtKY72  VKGDDVRQIAPAQTGVIADYNYKLPDDFTGCVLAWNTNSVDSKSGNN--FYYRLFRHGKIKPYERDISNVLYNSAGGTCSSISQ
pang17  VKGDEVRQIAPGQTGVIADYNYKLPDDFTGCVIAWNSVKQDALTGGNYGYLYRLFRKSKLKPFERDISTEIYQAGSTPCNGQVG
SARS2   IRGDEVRQIAPGQTGKIADYNYKLPDDFTGCVIAWNSNNLDSKVGGNYNYLYRLFRKSNLKPFERDISTEIYQAGSTPCNGVEG
RaTG13  ITGDEVRQIAPGQTGKIADYNYKLPDDFTGCVIAWNSKHIDAKEGGNFNYLYRLFRKANLKPFERDISTEIYQAGSKPCNGQTG
SHC014  VKGDDVRQIAPGQTGVIADYNYKLPDDFLGCVLAWNTNSKDSSTSGNYNYLYRWVRRSKLNPYERDLSNDIYSPGGQSCSA-VG
SARS1   VKGDDVRQIAPGQTGVIADYNYKLPDDFMGCVLAWNTRNIDATSTGNYNYKYRYLRHGKLRPFERDISNVPFSPDGKPCTP-PA
WIV1    VKGDDVRQIAPGQTGVIADYNYKLPDDFTGCVLAWNTRNIDATQTGNYNYKYRSLRHGKLRPFERDISNVPFSPDGKPCTP-PA
        :  .:***:**.:** **********:* ***:***:    *        : **  *  ::.*: **::.

                  500                 520                 540
Rs4081   -NGVRTLSTYDFYPNVPIEYQATRVVVLSFELLNAPATVCGPKLSTALVKNQCVNF
RmYN02   -NGVRTLSTYDFNPNVPLDYQATRVVVLSFELLNAPATVCGPKLSTQLVKNRCVNF
Rf1      -NGVRTLSTYDFNQNVPLEYQATRVVVLSFELLNAPATVCGPKLSTSLVKNQCVNF
BM-4831  LNCYKPLASYGFTQSSGIGFQPYRVVVLSFELLNAPATVCGPKQSTELVKNKCVNF
BtKY72   LGCYEPLKSYGFTPTVGVGYQPYRVVVLSFELLNAPATVCGPKKSTELVKNKCVNF
pang17   LNCYYPLERYGFHPTTGVNYQPFRVVVLSFELLNGPATVCGPKLSTTLVKDKCVNF
SARS2    FNCYFPLQSYGFQPTNGVGYQPYRVVVLSFELLHAPATVCGPKKSTNLVKNKCVNF
RaTG13   LNCYYPLYRYGFYPTDGVGHQPYRVVVLSFELLNAPATVCGPKKSTNLVKNKCVNF
SHC014   PNCYNPLRPYGFFTTAGVGHQPYRVVVLSFELLNAPATVCGPKLSTDLIKNQCVNF
SARS1    LNCYWPLNDYGFYTTTGIGYQPYRVVVLSFELLNAPATVCGPKLSTDLIKNQCVNF
WIV1     FNCYWPLNDYGFYITNGIGYQPYRVVVLSFELLNAPATVCGPKLSTDLIKNQCVNF
          .    *  *.*  .  : .*  **********:.******** ** *:*::****
```

* = fully conserved
: = strongly similar
. = weakly similar

**Supplementary Fig. 1. Sarbecovirus RBD sequence alignment**. Amino acid sequence alignment of sarbecovirus RBDs used in this study, numbered according to Spike protein of SARS2 Wuhan variant.

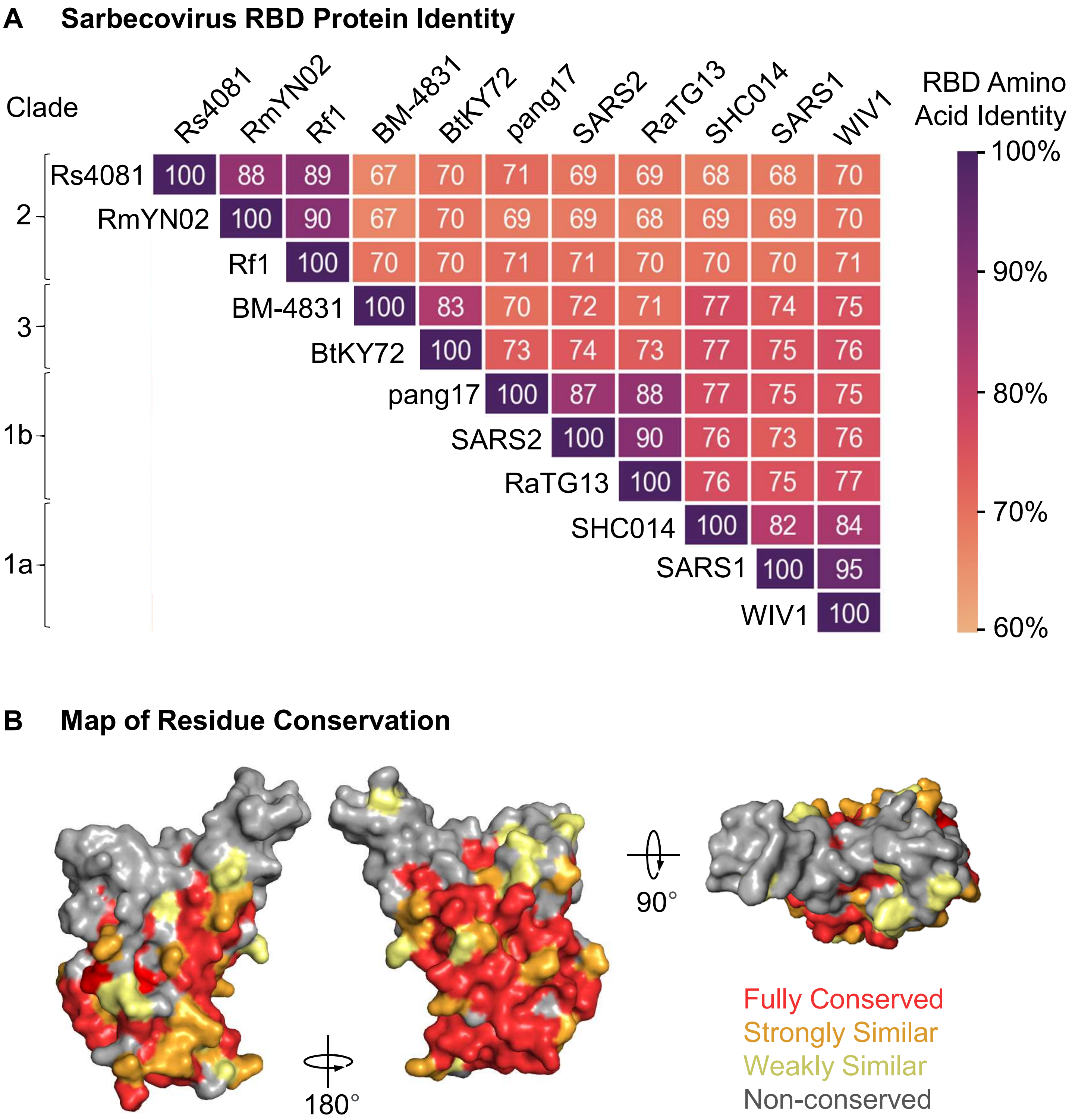


**Supplementary Fig. 2. RBD residue conservation.** (**A**) Heat map of percent amino acid identity between sarbecovirus RBDs used in this study. SARS2 refers to the Wuhan variant. (**B**) Conservation of residues between sarbecoviruses used in the study, as mapped onto the SARS2 RBD crystal structure (PDB ID: 6ZER). Multiple orientations of the same RBD are shown, represented as the van der Waals surface.

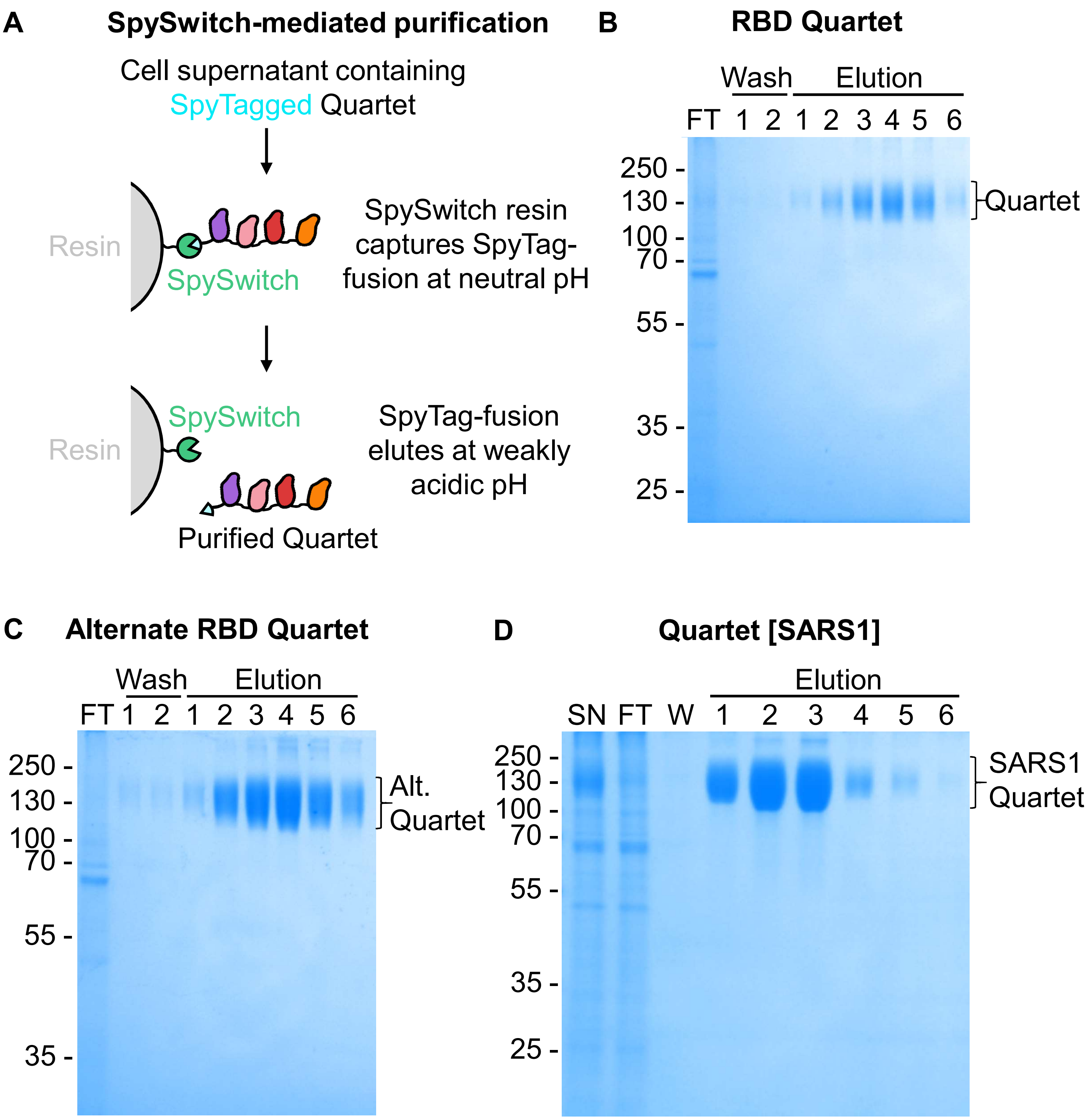


**Supplementary Fig. 3. SpySwitch purification of RBD quartets**. (**A**) Schematic of SpySwitch affinity purification. SpyTag genetically fused to the Quartet has a non-covalent interaction with SpySwitch at a neutral pH, before eluting at a weakly acidic pH through charge-charge repulsion. This system was used to purify (**B**) RBD Quartet, (**C**) Alternate RBD Quartet, and (**D**) RBD Quartet with SARS1 in place of SARS2. The supernatant (SN), flowthrough (FT), wash (W), and elution fractions were analyzed by SDS-PAGE with Coomassie staining.

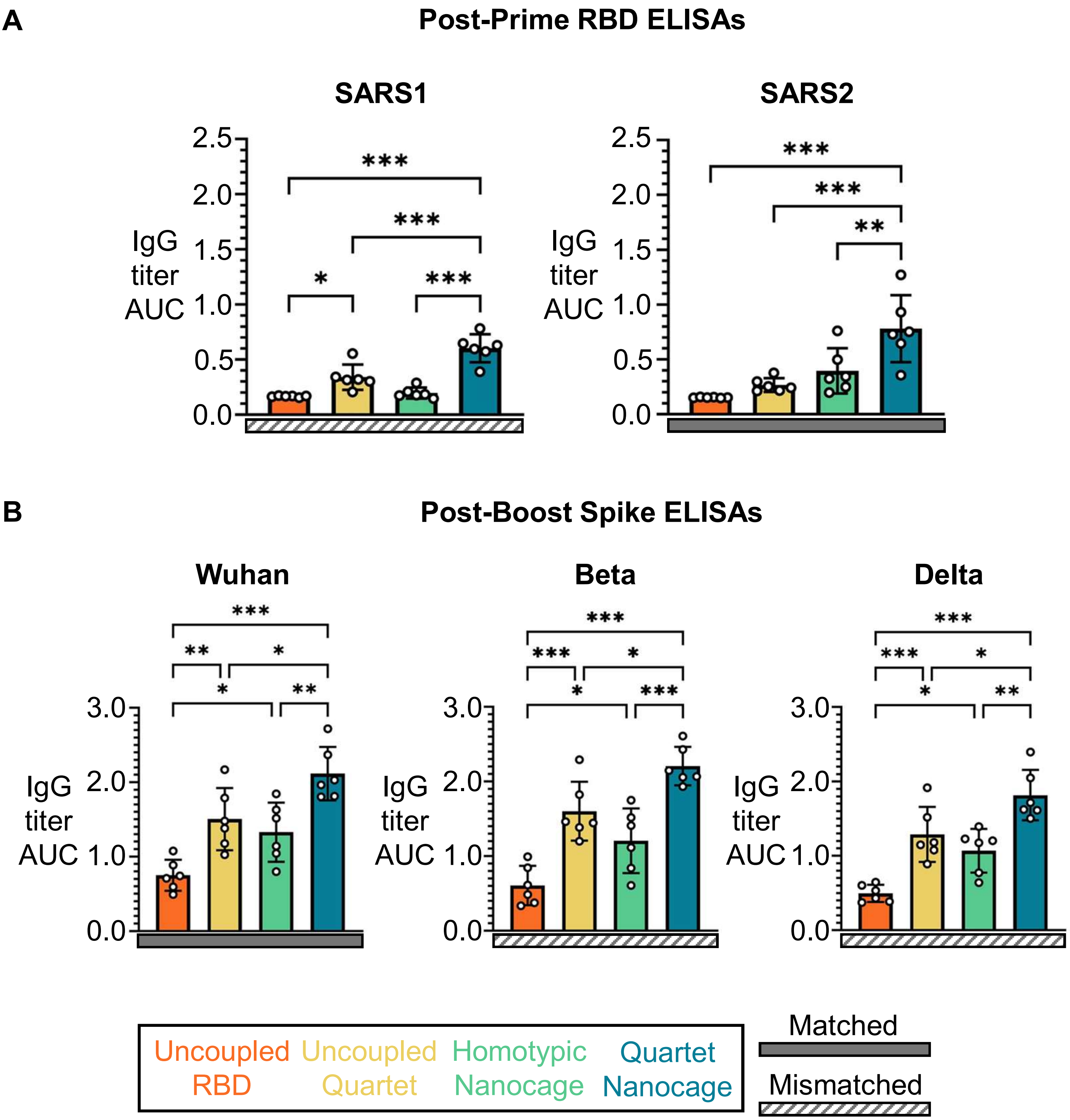


**Supplementary Fig. 4. Further Breadth of Immune Response from Immunization with Quartet Nanocages**. Binding data for serum IgG antibodies presented as area under the curve of a serial sera dilution. Sera samples are from mice immunized with uncoupled SARS2 RBD (orange), Uncoupled Quartet (yellow), SARS2 RBD coupled to SpyCatcher003-mi3 (Homotypic Nanocage, green), and Quartet Nanocage (blue) as outlined in Fig. 2. Solid gray rectangles under samples indicate the ELISA is against a component of that vaccine (matched), while striped rectangles indicate the ELISA is against an antigen absent in that vaccine (mismatched). Each dot represents sera from one animal. The mean is denoted by a bar, shown ± 1 s.d., n = 6. * $p < 0.05$, ** $p < 0.01$, *** $p < 0.001$; other comparisons were non-significant. Graphs demonstrate the binding of (**A**) post-prime sera to RBDs and (**B**) post-boost sera to SARS-CoV-2 variant Spike proteins.

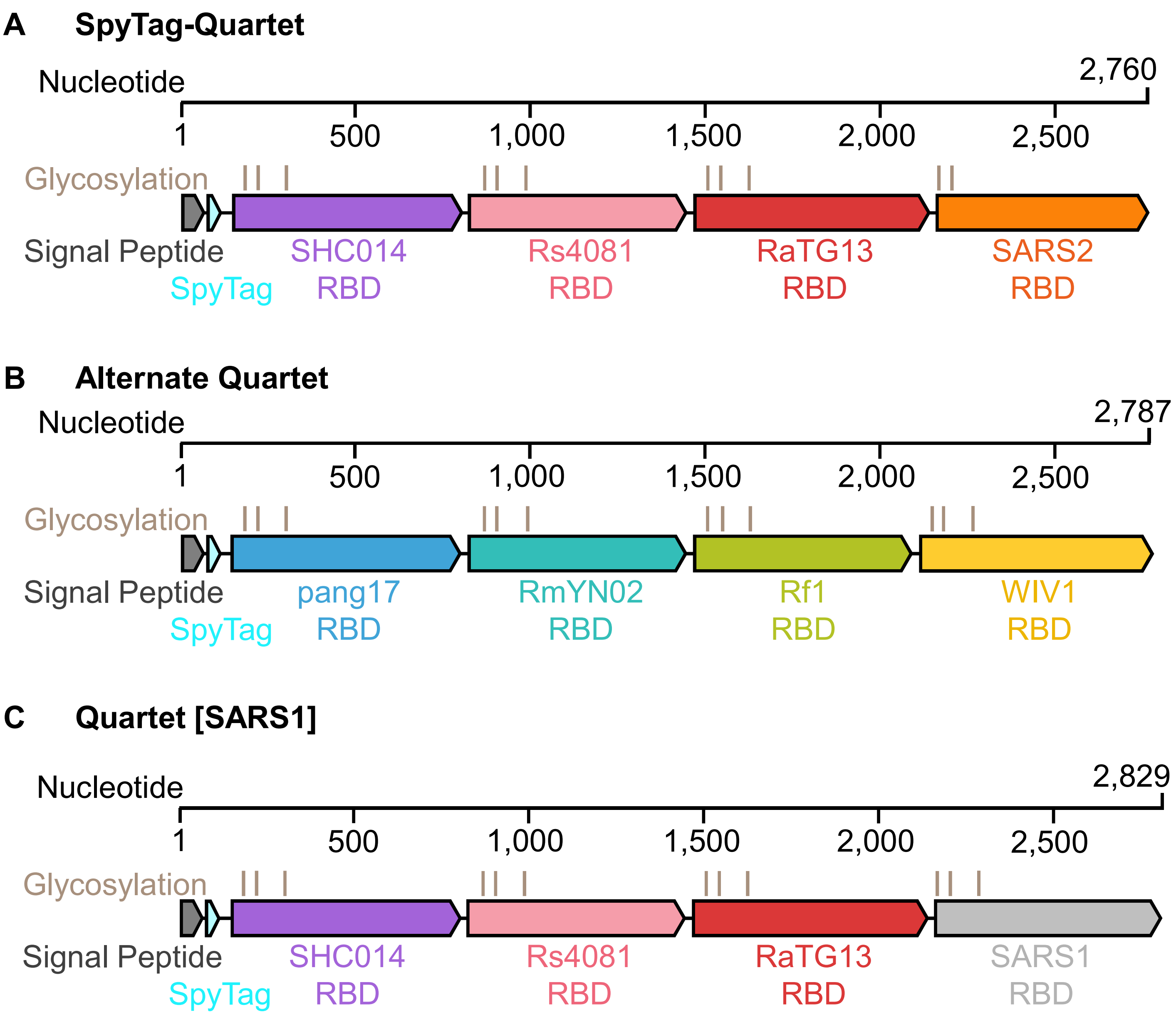


**Supplementary Fig. 5. Schematic of Different Quartets**. Genetic organization of (**A**) SpyTag-Quartet, (**B**) Alternate RBD Quartet, and (**C**) Quartet [SARS1]. These schematics indicate the virus origin of each RBD, predicted N-linked glycosylation sites, tag location, and nucleotide number for each construct.

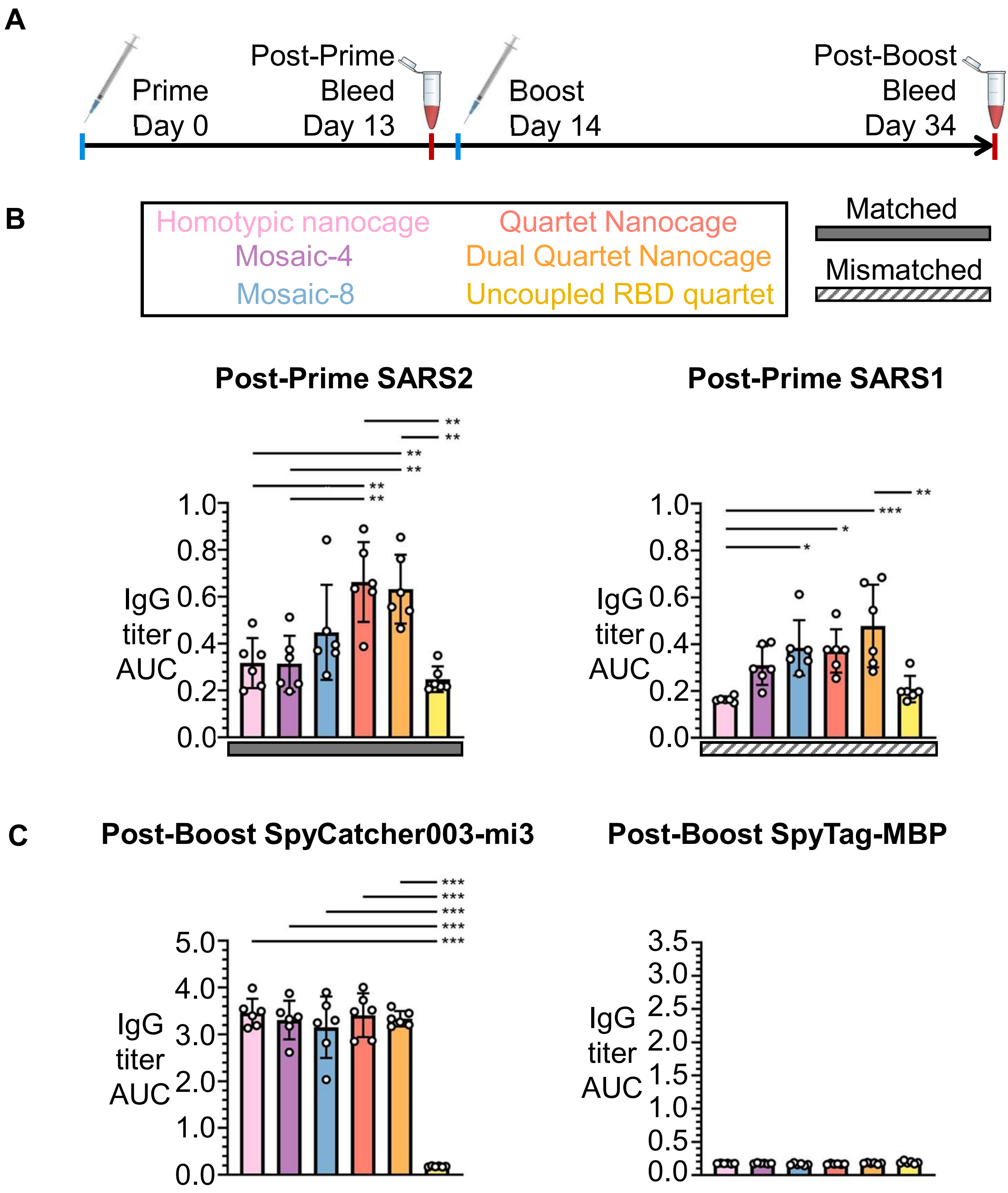


**Supplementary Fig. 6. Breadth of antibody induction by Quartet and Mosaic Immunogens.** (**A**) Summary of timeline for this set of immunizations with 0.02 nmol antigen per dose. **(B-C)** ELISA for serum IgG from mice immunized with the indicated immunogen. Each dot represents serum from one animal. The mean is denoted by a bar, with error bars ± 1 s.d., n = 6. * $p < 0.05$, ** $p < 0.01$, *** $p < 0.001$; other comparisons were non-significant. (**B**) Post-prime response to SARS2 or SARS1. (**C**) Post-boost response to SpyCatcher003-mi3 or SpyTag-MBP.

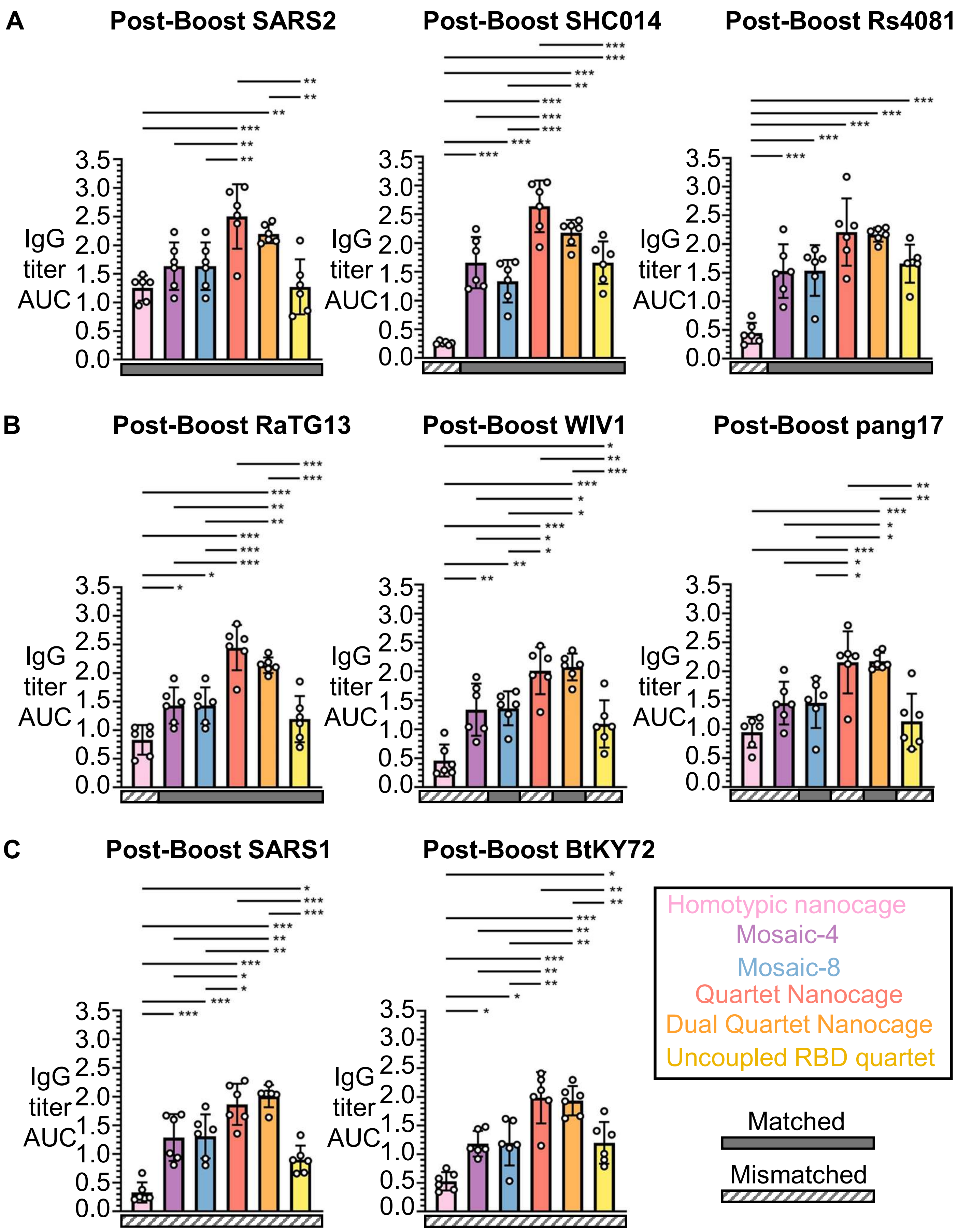


**Supplementary Fig. 7. Further breadth of antibody induction by Quartet and Mosaic immunogens.** ELISA for serum IgG from mice immunized with the indicated immunogen with 0.02 nmol antigen per dose. Each dot represents serum from one animal. The mean is denoted by a bar, with error bars ± 1 s.d., n = 6. * $p < 0.05$, ** $p < 0.01$, *** $p < 0.001$; other comparisons were non-significant. (**A**) Post-boost response to SARS2, SHC014 and Rs4081. (**B**) Post-boost response to RaTG13, WIV1 and pang17. (**C**) Post-boost response to SARS1 and BtKY72.

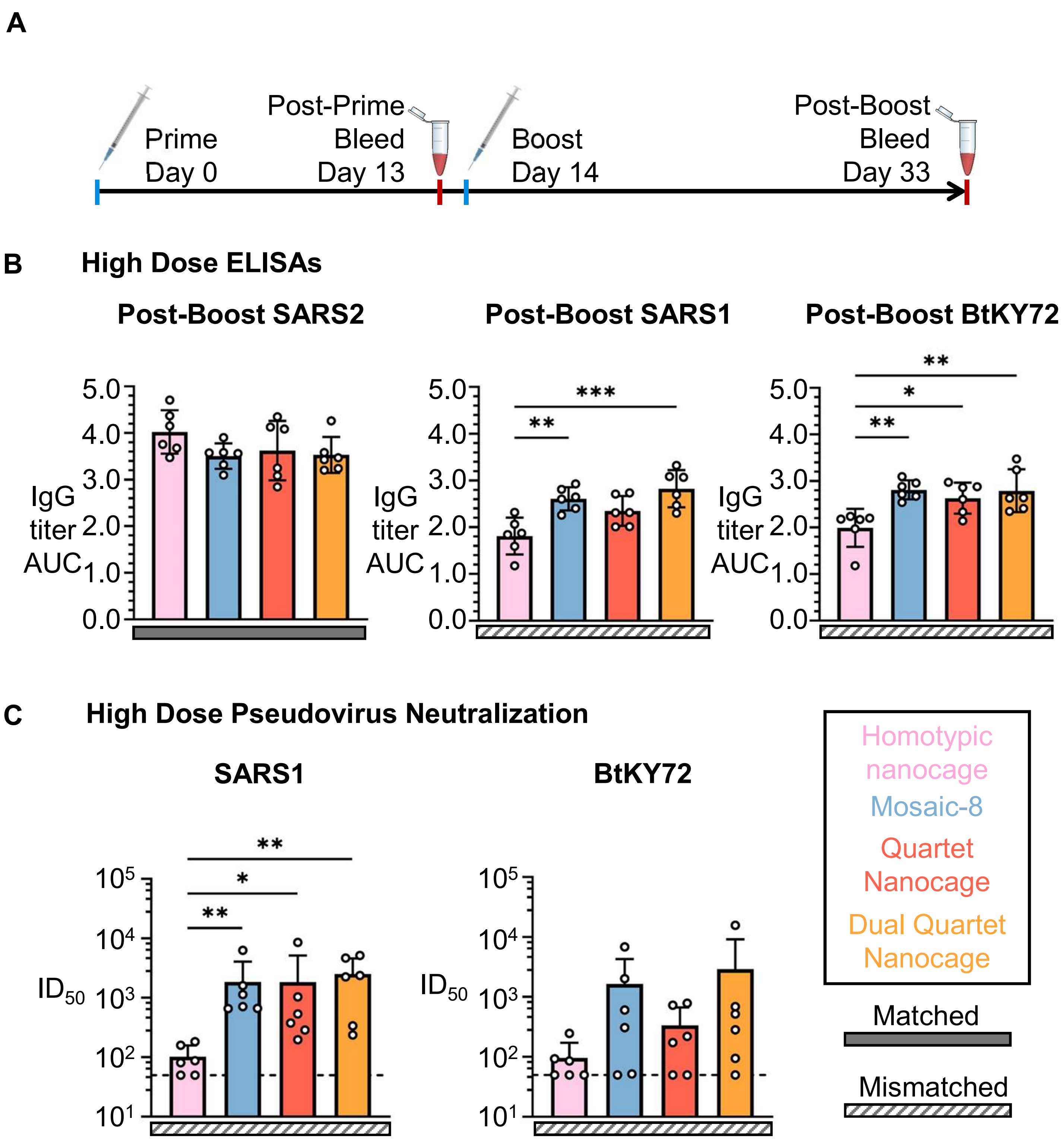


**Supplementary Fig. 8. Immune response raised by higher dose of Quartet and Mosaic immunogens.** This figure assesses antisera raised by immunizations with 0.2 nmol antigen. (**A**) Timeline for this set of immunizations. **(B)** ELISA for post-boost sera assessing IgG binding to SARS2, SARS1 and BtKY72 RBD is shown as the area under the curve (AUC) of a serial dilution. Each dot represents serum from one animal. The mean AUC is denoted by a bar, with error bars ± 1 s.d., n = 6. (**C**) Neutralization of SARS1 and BtKY72 (K493Y/T498W) pseudovirus by boosted mouse sera. Solid gray rectangles under samples indicate the ELISA is against a component of that vaccine (matched). Striped rectangles indicate the ELISA is against an antigen absent in that vaccine (mismatched). Dashed horizontal lines represent the limit of detection. The mean $ID_{50}$ is denoted by a bar, with error bars + 1 s.d., n = 6. * $p < 0.05$, ** $p < 0.01$, *** $p < 0.001$; other comparisons were non-significant.

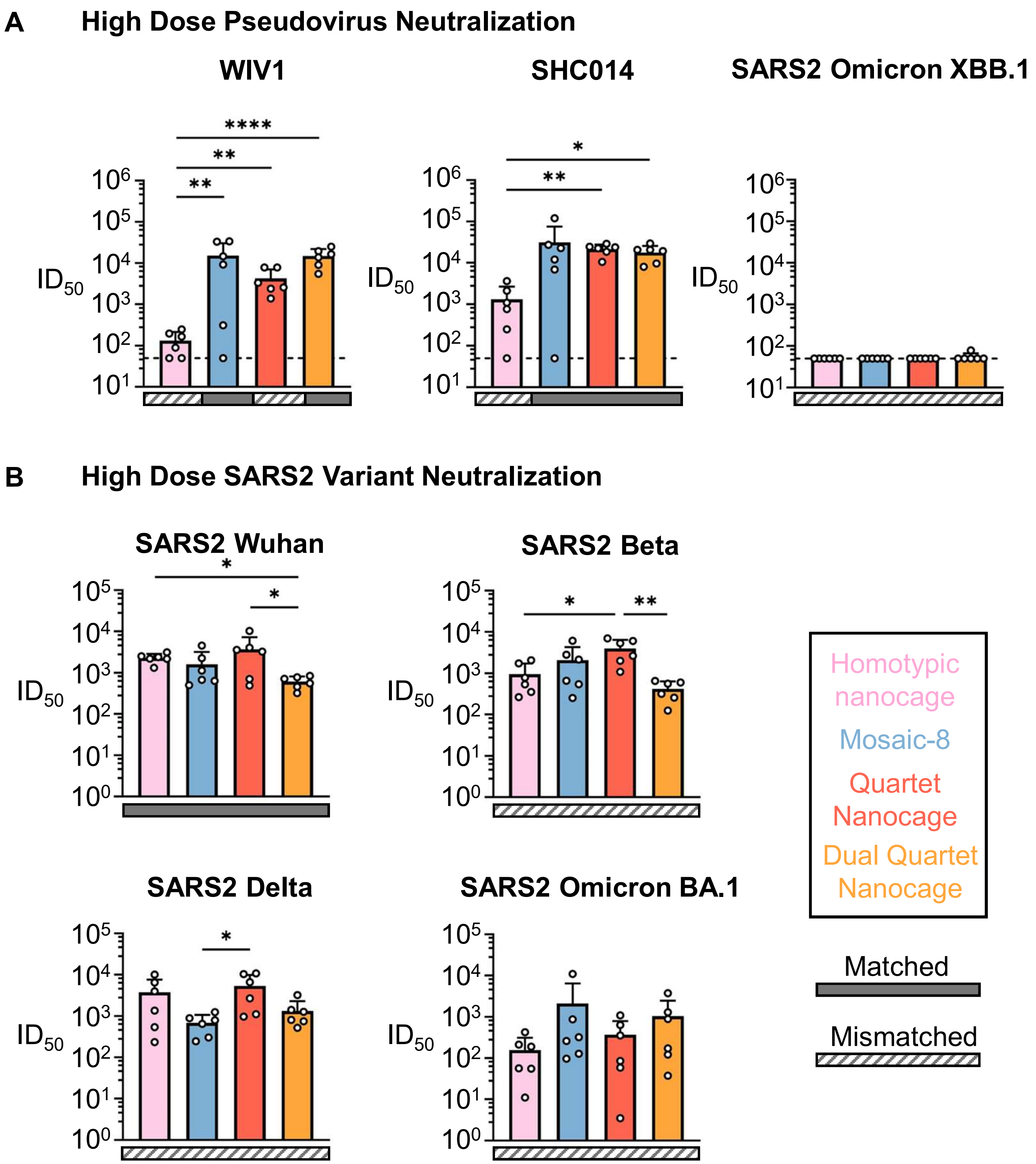


**Supplementary Fig. 9. Pseudovirus neutralization by higher dose of Quartet and Mosaic immunogens.** These figures assess antisera raised by immunizations with 0.2 nmol antigen, a 10x molar increase to antigen dose relative to prior immunizations. Solid gray rectangles under samples indicate the ELISA is against a component of that vaccine (matched). Striped rectangles indicate the ELISA is against an antigen absent in that vaccine (mismatched). Dashed horizontal lines represent the limit of detection. In all cases the mean $ID_{50}$ is denoted by a bar, with error bars + 1 s.d., n = 6. * $p < 0.05$, ** $p < 0.01$, *** $p < 0.001$; other comparisons were non-significant. (**A**) Neutralization of WIV1, SHC014 and SARS2 Omicron XBB.1 pseudoviruses. **(B)** Neutralization of the Wuhan, Beta, Delta and Omicron BA.1 SARS2 variant viruses.

### High Dose Virus Neutralization Curves

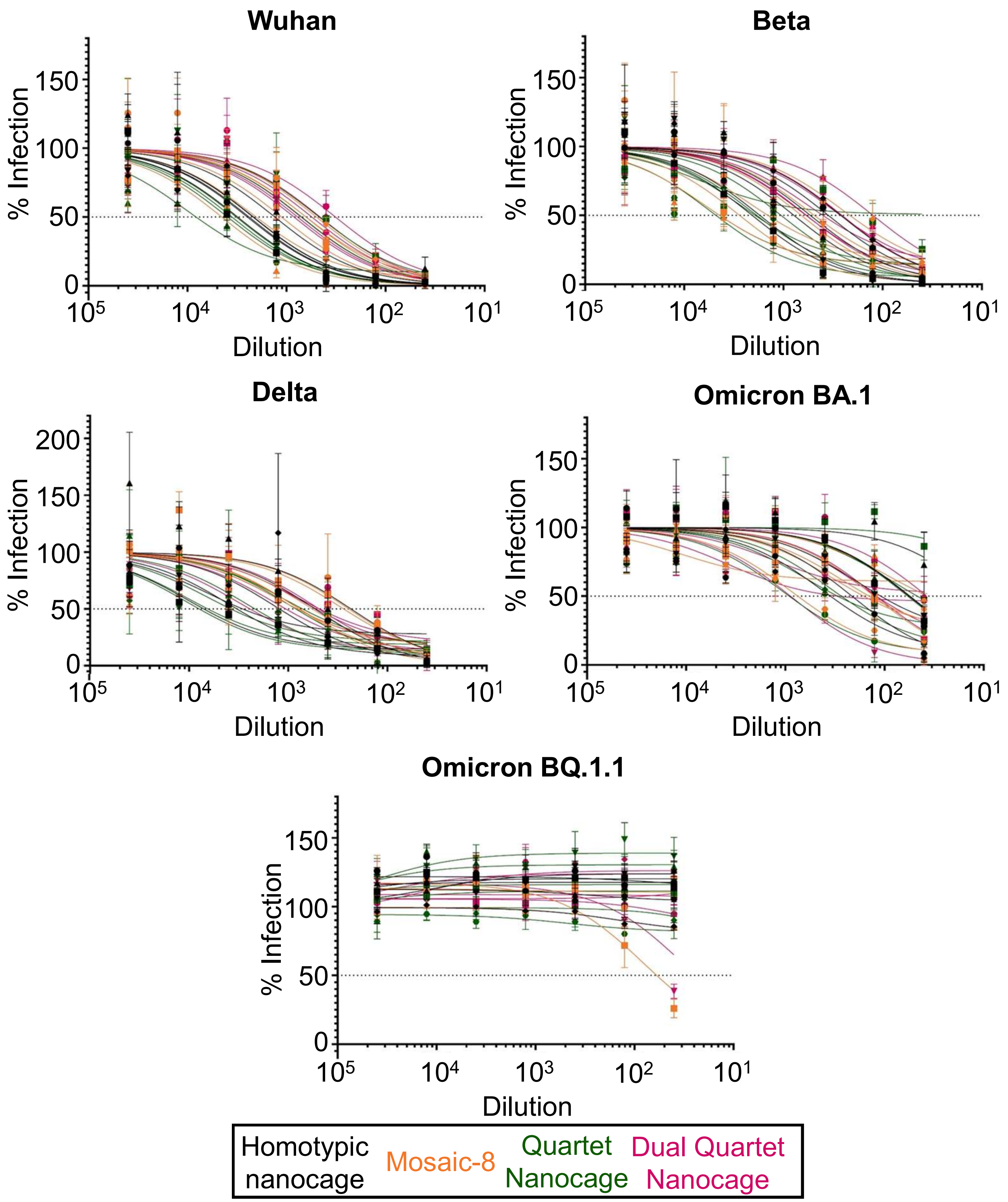


**Supplementary Fig. 10. Authentic virus dose response curves.** Neutralization of the Wuhan, Beta, Delta, Omicron BA.1, and Omicron BQ.1.1 SARS2 variants by antisera raised through immunizations with 0.2 nmol antigen. The percent of infection relative to a no-sera control (% Infection) was plotted relative to the dilution of sera. Each point is the mean of four replicates, with error bars ± 1 s.d., n = 6. These curves were used to determine the $ID_{50}$ values plotted in Supplementary Fig. 9B. $ID_{50}$ values could not be calculated for Omicron BQ.1.1.

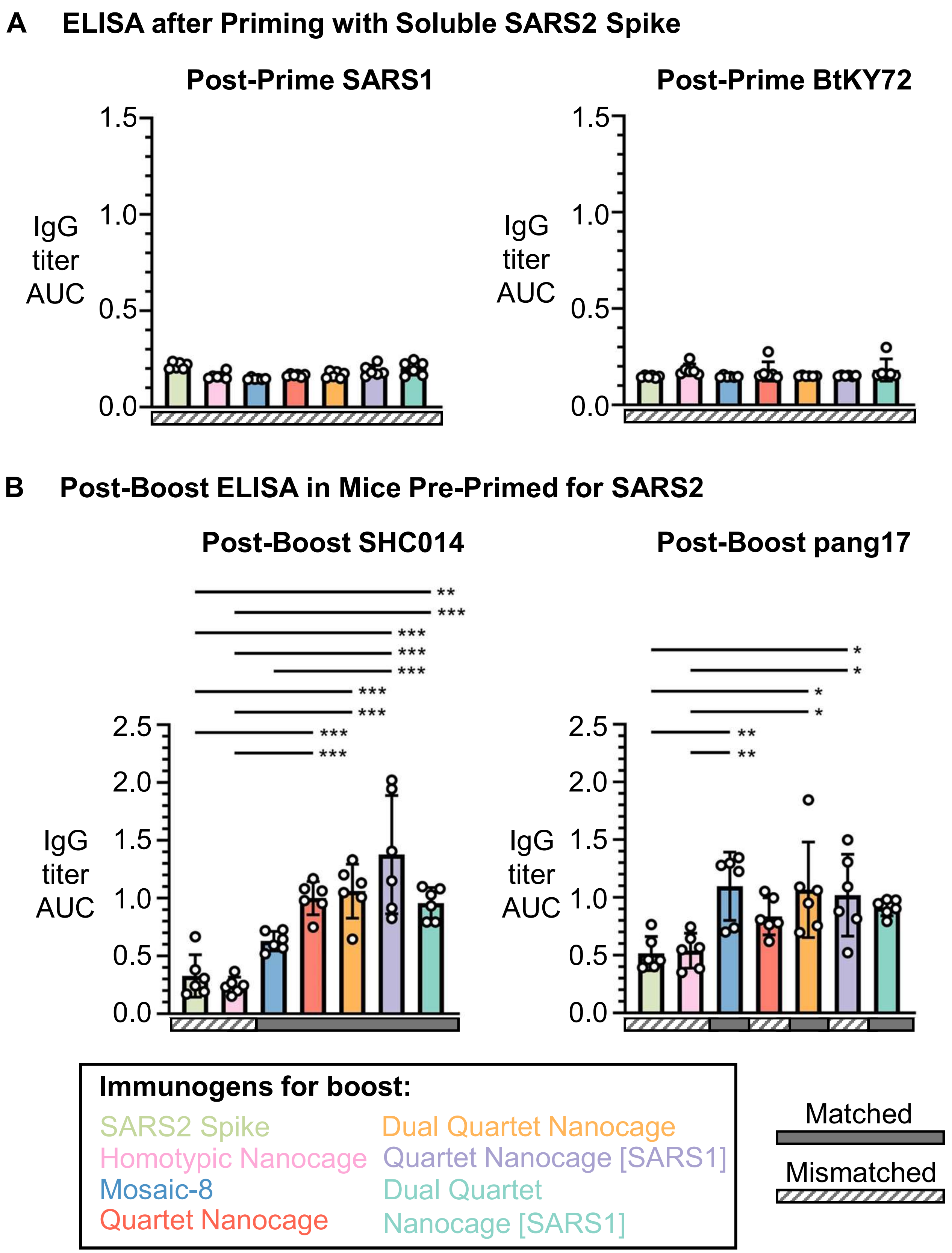


**Supplementary Fig. 11. Further demonstration that Quartet immunization induces broad antibodies even after SARS2 Spike priming.** (**A**) ELISA for serum IgG from mice immunized with a single dose of SARS2 Wuhan Spike protein, grouped by the second dose of 0.02 nmol antigen they will receive. (**B**) ELISA for serum IgG, after mice immunized with a single dose of SARS2 Wuhan Spike protein were boosted with a variety of different antigens at 0.02 nmol per dose. Each dot represents serum from one animal. The mean is denoted by a bar, with error bars ± 1 s.d., n = 6. * $p < 0.05$, ** $p < 0.01$, *** $p < 0.001$; other comparisons were non-significant.
